# Rapid evolution of enhanced Zika virus virulence during direct vertebrate transmission chains

**DOI:** 10.1101/2020.10.23.353003

**Authors:** Kasen K. Riemersma, Anna S. Jaeger, Chelsea M. Crooks, Katarina M. Braun, James Weger-Lucarelli, Gregory D. Ebel, Thomas C. Friedrich, Matthew T. Aliota

## Abstract

Zika virus (ZIKV) has the unusual capacity to circumvent natural alternating mosquito-human transmission and be directly transmitted human-to-human via sexual and vertical routes. The impact of direct transmission on ZIKV evolution and adaptation to vertebrate hosts is unknown. Here we show that molecularly barcoded ZIKV rapidly adapted to a mammalian host during direct transmission chains in mice, coincident with the emergence of an amino acid substitution previously shown to enhance virulence. In contrast, little to no adaptation of ZIKV to mice was observed following chains of direct transmission in mosquitoes or alternating host transmission. Detailed genetic analyses revealed that ZIKV evolution in mice was generally more convergent and subjected to more relaxed purifying selection than in mosquitoes or alternate passages. These findings suggest that prevention of direct human transmission chains may be paramount to resist gains in ZIKV virulence.

## Introduction

Following its discovery in central Africa in 1947, Zika virus (ZIKV; genus *Flavivirus*, family *Flaviviridae*) was introduced and circulated in Asia unbeknownst to the scientific community (Petersen et al., 2016). Outbreaks in Yap in 2007 and French Polynesia in 2013 established ZIKV as a pathogen of global concern. Those concerns were realized in 2015 and 2016, when Asian-lineage ZIKV swept through nearly every country in the Americas (Hills et al., 2017; Petersen et al., 2016). The reasons for the sudden emergence of ZIKV in the Americas are still not completely understood. As a result, extensive research efforts have aimed to understand whether there are mutations that have contributed to the severity of the ZIKV outbreak in the Americas. Mounting evidence indicates that African-lineage and American-sublineage ZIKV are generally more transmissible than other Asian lineage ZIKVs (Aubry et al., 2020; Calvez et al., 2018; Liu et al., 2017; Pompon et al., 2017; Roundy et al., 2017; Weger-Lucarelli et al., 2016). Further, a panel of mutations present in American-sublineage ZIKV, but not other Asian-lineage ZIKV, were shown to enhance transmission by *Aedes aegypti* mosquitoes, as well as replication and virulence in mouse models (Liu et al., 2020; Shan et al., 2020). While there are data on the effects of single mutations in ZIKV, there is a need to better understand the biological basis of mutational effects.

ZIKV is a mosquito-borne virus that naturally cycles between vertebrate hosts and mosquito vectors. In urban environments, transmission predominantly occurs between humans and *Ae. aegypti* mosquitoes. In both field and laboratory settings, female *Ae. aegypti* have been shown to transmit ZIKV to their progeny at low rates, indicating that the virus can bypass the vertebrate host (Ciota et al., 2017; Comeau et al., 2020; da Costa et al., 2018; Thangamani et al., 2016). Similarly, ZIKV can bypass the mosquito vector with direct human-to-human transmission. Of great clinical concern, vertical transmission during pregnancy can result in congenital Zika syndrome, a term for the combination of neuropathologic birth defects and disabilities following *in utero* exposure to ZIKV (Cristina da Silva Rosa et al., 2020; Gregory et al., 2017; Grischott et al., 2016; Moore et al., 2017). ZIKV can also be transmitted horizontally in humans via sexual intercourse (Foy et al., 2011; Fréour et al., 2016; Hills et al., 2016; McCarthy, 2016; Moreira et al., 2017; Turmel et al., 2016; Venturi et al., 2016). Rare cases of horizontal transmission via blood transfusion (Barjas-Castro et al., 2016), breastfeeding (Colt et al., 2017), and nonsexual contact (Swaminathan et al., 2016) have also been reported. Rates of sexual ZIKV transmission and its contribution to epidemic spread are difficult to quantify in regions with endemic *Ae. aegypti*, but the increased risk for seropositivity in sexual partners of index cases (Magalhaes et al., 2020) and secondary cases in *Ae. aegypti*-free regions (Grischott et al., 2016; Khaiboullina et al., 2019) suggest it is a common non-vector-borne route of transmission. Given the aforementioned evidence for ZIKV mutations enhancing transmission and virulence, it is critical to study the impact of bypassing the mosquito vector on ZIKV evolution and the potential for adaptation to vertebrate hosts.

For arthropod-borne viruses (arboviruses), it has been proposed that a fitness trade-off occurs during host cycling, where fitness gains in one host are counteracted by fitness losses in the opposing host (Ciota and Kramer, 2010). However, this has not been supported by *in-vivo* infection studies generally (Ciota et al., 2009, 2008; Coffey et al., 2008; Deardorff et al., 2011). Release from host cycling does typically enable adaptation to the vertebrate or arthropod, but not necessarily at the cost of lost fitness in the other host. We hypothesized that ZIKV would demonstrate a similar capacity for host adaptation when bypassing the mosquito vector, an outcome with potentially significant implications during direct ZIKV transmission in humans. Here, we tested this hypothesis by assessing phenotypic and genotypic changes following serial passage in mice or mosquitoes, and during alternating passage between both, using a molecularly barcoded ZIKV strain previously validated for tracking genetic bottlenecks and selective pressures within mosquitoes and non-human primates (Aliota et al., 2018; Weger-Lucarelli et al., 2018). We found that ZIKV rapidly acquires enhanced virulence with universal fatality in mice coincidental to selective sweeps involving a previously described virulence-enhancing mutation. We additionally show that ZIKV populations evolve convergently under relaxed purifying selection in vertebrate hosts, whereas stochasticity and purifying selection characterize ZIKV evolution in mosquitoes and during alternating transmission.

## Results

### Conspecific and alternating passage titers

To determine the effect of release from host-cycling on ZIKV evolution, *in-vivo* serial passage experiments were conducted with a previously characterized barcoded ZIKV (ZIKV-BC; strain PRVABC59) containing a run of eight consecutive degenerate codons in NS2A (amino acids 144-151; Figure 1A) that allows for every synonymous mutation to occur. There was no evidence of barcode bias in our ZIKV-BC stocks, as the 8,811 barcodes detected by deep sequencing were evenly distributed at less than 0.3% frequency within the population (Figure 1B). ZIKV-BC was serially passaged in 5 parallel replicates (lineages) for 10 passages via subcutaneous (SQ) inoculation in *Ifnar1*^−/−^ mice or via intrathoracic (IT) inoculation in *Ae. aegypti* mosquitoes (Figure 1C). Despite a consistent amount of infectious virus inoculated at each passage, ZIKV-BC titers rose significantly over 10 serial passages in both mice and mosquitoes (non-zero slope F-test, *P*<0.0001; Figure 2A). The mean increase in infectious virus titer from passage 1 to 10 was greater in mice (+ 10^2.97^ pfu/ml) than in mosquitoes (+ 10^0.15^ pfu/ml).

**Figure 1.**
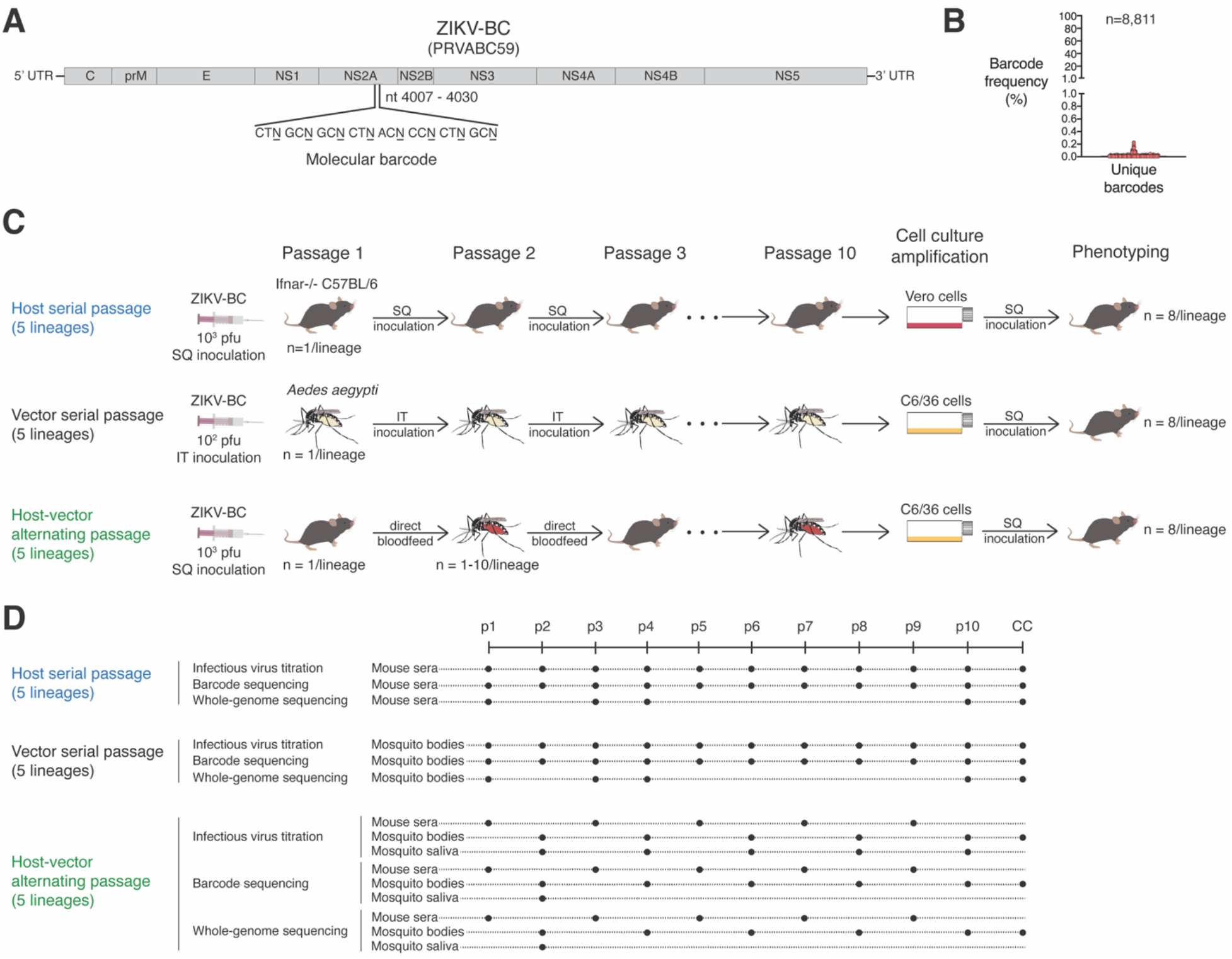
Genetic and phenotypic evolution of molecularly barcoded ZIKV tracked during serial or alternating *in vivo* passage. **(A)** Stocks of molecularly barcoded ZIKV (ZIKV-BC) consist of **(B)** an unbiased distribution of unique barcodes. **(C)** Five replicate lineages of ZIKV-BC were serially passaged via needle inoculation for ten passages in Ifnar1^−/−^ C57BL/6 mice or *Aedes aegypti* mosquitoes, or alternately passaged via bloodfeeding for ten passages. Passage 10 viruses were amplified once in Vero or C6/36 cells before phenotypic analysis in mice. **(D)** Viral replication was tracked by plaque assay after each passage. Deep sequencing of virus barcodes and whole ZIKV genomes was employed to characterize virus population structure and composition, respectively, over the course of ten passages. SQ: subcutaneous, IT: intrathoracic, and CC: cell culture.

**Figure 2.**
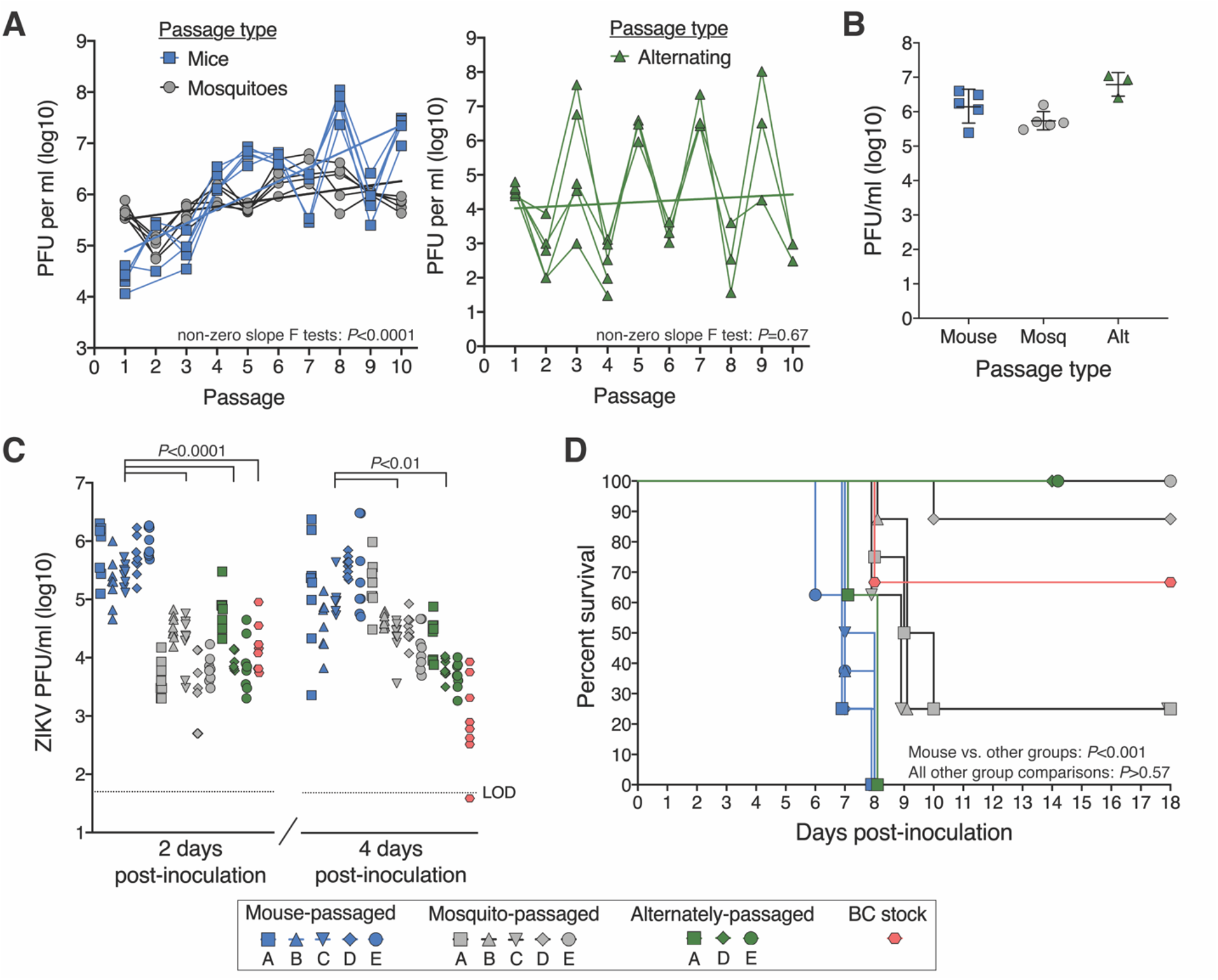
Phenotypic changes of serially and alternately passaged ZIKV-BC in mice. **(A)** Infectious ZIKV titers over sequential serial or alternating passages. **(B)** High titers of cell culture-amplified passage 10 viruses. **(C)** Infectious titers of passage 10 viruses and ZIKV-BC stock in mouse sera 2 and 4 days post-inoculation. *P*-values shown are from ANOVA tests for each day post-inoculation. **(D)** Survival of mice inoculated with passage 10 viruses and ZIKV-BC stock. *P*-values shown are from Mantel-Cox log-rank tests between each group. PFU: plaque-forming unit, Alt: alternate, LOD: limit of detection. Symbol shapes and letters represent replicate lineages.

To mimic natural host-cycling, ZIKV-BC was alternately passaged in mice and mosquitoes for 10 passages in 5 parallel lineages. After SQ inoculation of mice on passage 1, alternating passage was conducted via natural bloodfeeding transmission with small cohorts of mosquitoes feeding on an infected mouse, and then feeding on a naive mouse 12 days later. Only 3 of 5 lineages successfully completed the series of 10 passages. Despite successful bloodfeeding, lineage B and C viruses replicated to very low titers (<10^2^ pfu/ml) in mosquito bodies on passage 4 and were not transmitted onwards to mice. Unlike with serial conspecific passage, ZIKV-BC titers did not rise over the course of 10 alternating passages (non-zero slope F-test, *P*=0.67). The rising and falling sequential titers with alternating passage reflect the virus’ capacity for greater replication in mice than mosquitoes. After passaging ZIKV-BC with and without host-cycling, we next assessed the p10 viruses for phenotypic changes in viral replication and virulence in mice.

### Phenotypic changes in mice

Viral replication and virulence of mouse-adapted, mosquito-adapted, and alternately-passaged p10 lineages were compared with unpassaged ZIKV-BC stock in *Ifnar1*^−/−^ mice. To generate adequate viral stocks of the passaged viruses, each p10 virus isolate was amplified in a single passage at high multiplicity of infection on Vero cells for mouse-adapted lineages or on a mosquito cell line (C6/36) for mosquito-adapted and alternately-passaged lineages (Figure 2B). Cell culture-amplified stocks and p10 virus isolates were deep sequenced to ensure minimal changes in the single nucleotide variant (SNV) frequencies after amplification (Figure 2 — supplemental figure 1). To evaluate viral replication, infectious virus was titered by plaque assay at 2 and 4 days post-inoculation (dpi) from mouse sera (Figure 2C). At 2 dpi, only mouse-adapted ZIKV replicated to significantly higher titers compared to the unpassaged ZIKV-BC (one-way ANOVA, *P*<0.0001). By 4 dpi, both mouse-adapted and mosquito-adapted lineages replicated to higher titers compared to the unpassaged ZIKV-BC (one-way ANOVA, *P*<0.0001). In terms of virulence, only the mouse-adapted lineages were associated with reduced survival in mice compared to unpassaged virus (Log-rank tests, *P*=0.0009; Figure 2D). Median survival was 7 dpi for mice infected with the mouse-adapted lineages with all mice succumbing by 8 dpi. The mosquito-adapted and alternately-passaged lineages demonstrated a wide range of virulence, with some lineages producing enhanced mortality rates relative to unpassaged virus, and other lineages generating little to no mortality. These data suggest that focused adaptation to a specific environment broadened diversity with several phenotypic variants arising, suggesting that host specialization can alter the direction of virulence evolution in virus populations. We therefore aimed next to define genotypic diversity and the impact on ZIKV population structure/composition associated with differences in virus replication and virulence.

### Barcode dynamics

We deep sequenced the barcode in virus populations from our passage series, which allowed us to track changes in the ZIKV population structure over sequential passages. In serial mouse and serial mosquito lineages, unique molecular identifiers (UMI) were incorporated in the barcode sequencing libraries to enable bioinformatic filtering of PCR and sequencer errors that could create false barcodes. The UMI approach was not possible with the mosquito samples (even-numbered passages) in the alternating passage lineages due to low concentrations of ZIKV RNA. Instead, whole genome sequencing reads covering the entire barcode region were processed with stringent quality-trimming and overlapping-read error correction, and then analyzed in the same manner as the UMI approach. For alternating passages, only mosquitoes with detectable infectious virus in their body and saliva were sequenced. In alternating passage lineages C and E, the dominant barcode in passage 3 and onwards was not detected in any of the sequenced mosquito tissues from passage 2, indicating that onward transmission was instigated by a mosquito whose saliva was not sequenced. As a result, mosquito bodies and saliva at passage 2 in lineages C and E were excluded from barcode analyses and figures.

Across the five lineages serially passaged in mice, the virus populations were composed of more than 10^3^ uniquely barcoded viruses until passages 3 and 4, after which the populations were rapidly overtaken by a small number of barcoded viruses that remained dominant through passage 10 (Figure 3A-B, Figure 3 — supplemental figure 1A). In contrast, the virus populations serially passaged in mosquitoes exhibited a slower and steadier loss of population structure heterogeneity from approximately 10^3^ uniquely barcoded viruses to 10^2^ over the ten passages (Figure 3A-B, Figure 3 — supplemental figure 1B). The divergence in the viral barcode populations relative to passage 1 was measured by Euclidean distance. Despite achieving comparable divergence by passage 10, the dynamics of genetic divergence in barcode populations differed significantly between serial mouse lineages and serial mosquito lineages over ten passages (Wilcoxon matched-pairs signed rank test, *P*=0.03; Figure 3C). During serial mouse passage, rapid divergence in the first four passages was followed by slower divergence in the final six passages, whereas the rate of divergence was relatively stable during serial mosquito passage. For the alternately passaged populations, the population structure contracted down to only a few unique barcodes after the first passage in mosquitoes (Figure 3A,B, Figure 3 — supplemental figure 1C). In line with known anatomical bottlenecks in the mosquito midgut and salivary glands, constriction of the population structure was observed in mosquito bodies with further constriction in the mosquito saliva (Figure 3A). Sudden homogenization of virus population structure, as seen during serial mouse and alternate passaging, is consistent with either a stringent genetic bottleneck or selective sweep(s).

**Figure 3.**
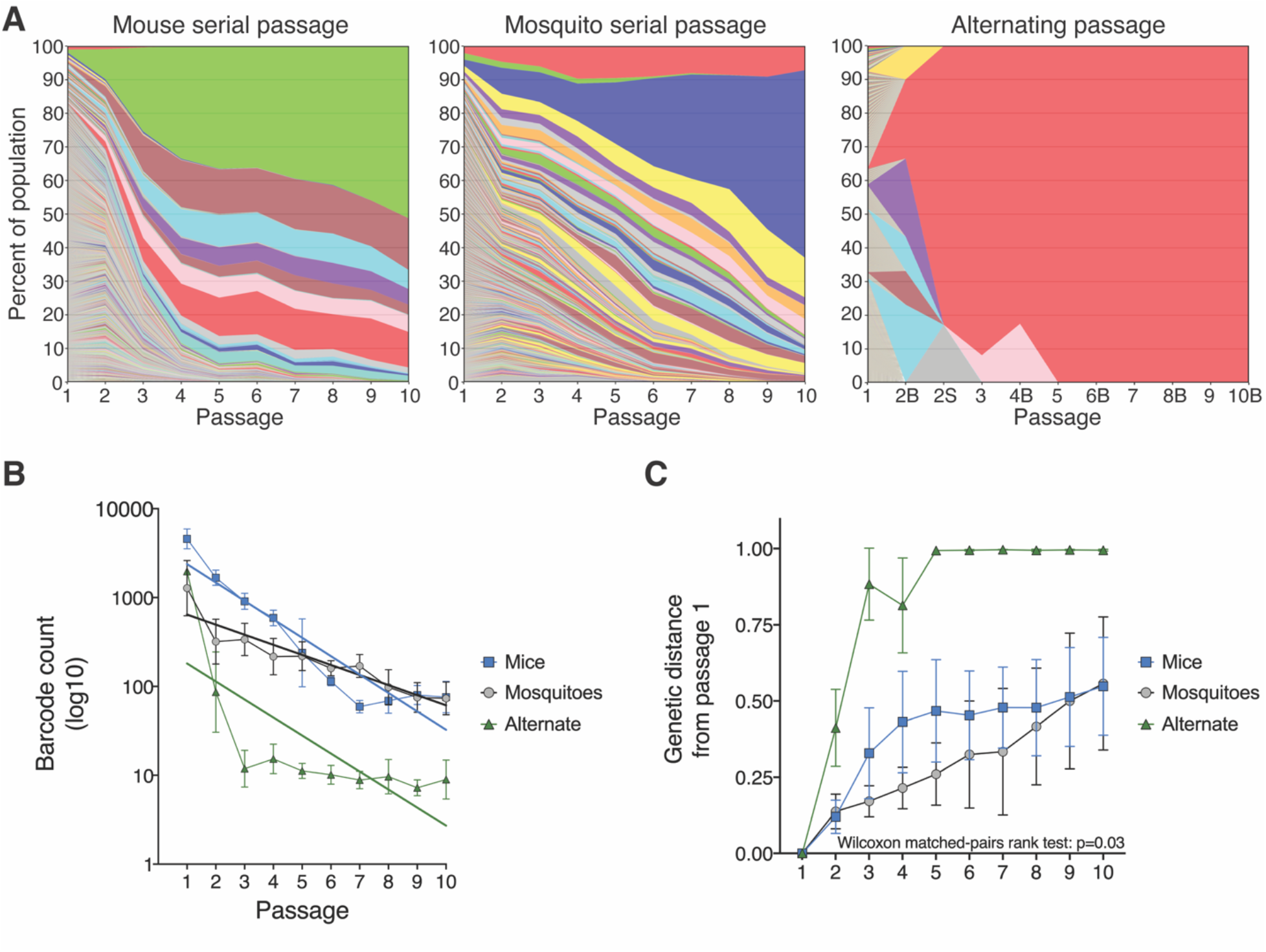
ZIKV barcode dynamics over sequential passages. **(A)** Individual barcode frequencies over 10 serial or alternating passages. Composite images for each passage series were generated by ranking barcodes from most to least frequent and calculating the mean frequency of the barcodes at each rank across the five replicate lineages. Colors represent barcode rank and are not associated with the barcode sequence. Thus, each colored bar is the mean frequency of barcodes with the same rank in the five replicate lineages. In the alternating passage series, even passages are labeled with “B” or “S” to differentiate mosquito bodies and saliva, respectively. **(B)** Barcode abundance over sequential passages. Solid lines are linear regression lines of best fit. **(C)** Euclidean distance of barcode populations relative to populations at passage 1. Values ranging from 0 to 1 indicate degree of genetic similarity, with lower values indicative of high similarity and vice versa. Wilcoxon matched-pairs rank test result reported for comparison between serial mouse and serial mosquito passage.

Importantly, none of the 50 most frequent barcodes present at passage 10 were shared between any mosquito lineage or alternate lineage (Figure 3 — supplemental figure 2A), indicating phenotypic neutrality for the barcodes *in vivo*. Of the 50 most frequent barcodes present at passage 10 in mouse lineages, ten were present in more than one lineage, although there was no positive correlation between barcode frequencies across lineages (Spearman’s correlation, *P*=0.51; Figure 3 — supplemental figure 2B). While selective advantages for certain barcodes in mouse cannot be ruled out, early selective sweeps and genetic hitchhiking can also cause barcode sharing if the barcode is linked with a non-barcode mutation in the virus stocks that is selected for *in vivo*. We therefore conclude that the genetic barcodes are unbiased, neutral reporters of population evolution. In addition to evaluating ZIKV population structure, we also performed whole genome deep sequencing to assess changes in the genetic composition of the ZIKV population.

### ZIKV host-adaptation associated with consistent emergence of NS2A A117V

Informed by the observed population structure dynamics in mice, we selected passages 1, 3, 4, 10, and the cell culture amplification passage of the serial mouse and mosquito passage lineages for whole-genome deep sequencing. For the alternating transmission lineages, all 10 passages and the cell culture amplification passage were sequenced (Supplemental Table 1). The frequency of individual SNVs were tracked over sequential passages to monitor the dynamics of ZIKV population composition. All SNVs called at greater than 1% frequency in any passage and with at least 300 reads of coverage were tracked. Depth of coverage was greater than 300 reads across the entire coding region for 81% (165/204) of sequencing libraries, with high coverage on at least 70% of the coding region in the remaining libraries (Figure 4 — supplemental figure 1). SNVs were further compared across lineages and passage series to assess convergent evolution. As with barcode sequencing, passage 2 from alternating passage lineages C and E was excluded from SNV analyses and figures since the mosquito contributing to onward transmission was not sequenced. For the alternate passage lineages A, B, and D, passage 2 SNV data are from the single mosquito that contributed to onward transmission. SNV data from passages 4, 6, 8, and 10 for all alternate passage lineages are from mosquito pools since the mosquito(es) contributing to onward transmission could no longer be identified by barcode sequences.

**Figure 4.**
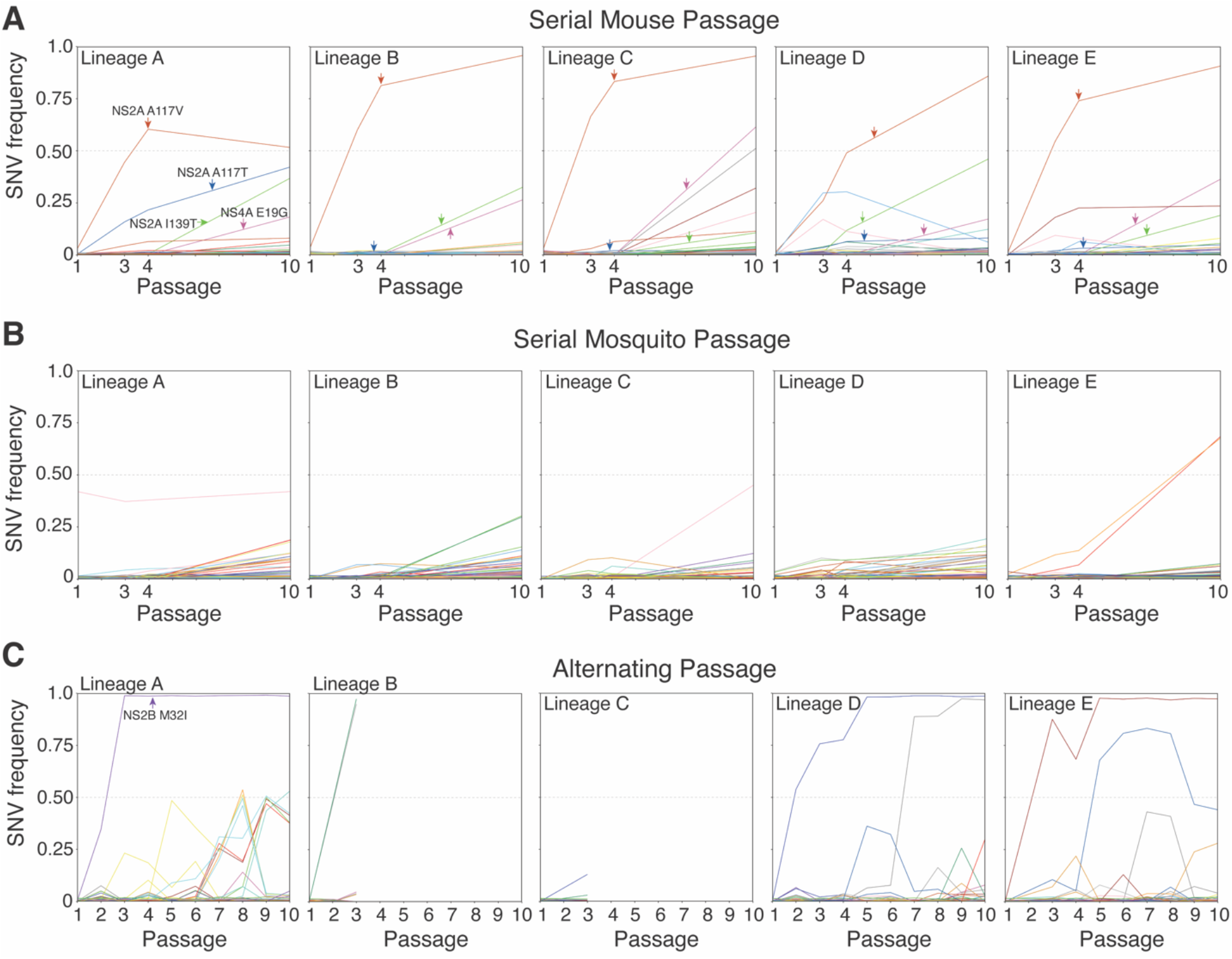
Trajectories of individual single-nucleotide variants (SNVs) over sequential passages detected at >1% frequency in any passage. SNV frequencies for individual SNVs detected in. **(A)** serial mouse lineages, **(B)** serial mosquito lineages, **(C)** alternating passage lineages. Four non-synonymous SNVs detected in all five mouse lineages are demarcated with arrows, and one SNV of note is similarly demarcated in alternating passage lineage A. Colors represent the same SNVs across homotypic lineages, but are used more than once due to the large number of SNVs. Also, the same SNVs may not be represented by the same colors across passage types. In the alternating passage series, odd passages are mouse sera and even passages are mosquito bodies.

In the mouse serial passage lineages, four non-synonymous SNVs, NS2A A117V, NS2A A117T, NS2A I139T, and NS4A E19G, arose in all five lineages, typically reaching high frequency in the population (Figure 4A). Of particular note, the NS2A A117V rose from less than 2% frequency at passage 1 to greater than 25% by passage 3 and greater than 45% by passage 4. In four of the five lineages, NS2A A117V was present on more than 75% of viruses at passage 10. Interestingly, in lineage A, the frequency of NS2A A117V plateaued at just above 50%, but another mutation at the same locus, NS2A A117T, rose to 40% by passage 10, such that in this lineage viruses encoding alternate amino acids at NS2A residue 117 accounted for more than 90% of the population. The less frequent NS2A A117T mutation was also found at low frequency (<10%) in the other four lineages. The trajectory of the NS2A polymorphisms aligns closely with the aforementioned population structure dynamics (Figure 3C), supporting our hypothesis that selective sweeps accounted for the dramatic loss in viral population diversity during mouse passage. The two other SNVs found in all five lineages, NS2A I139T and NS4A E19G, tended to arise between passages 4 and 10, and were typically found on less than 50% of the ZIKV genomes. None of the four presumed mouse-adaptive mutations demonstrated parallel trajectories indicative of genetic hitchhiking. The two NS2A 117 mutations and NS4A E19G were not detected in any passage of the serial mosquito and alternating passages. The NS2A I139T was detected in two serial mosquito lineages, but never at greater than 2% frequency.

Unlike with serial mouse passage, we detected no SNVs shared by all 5 serial mosquito lineages above our 1% frequency cut-off (Figure 4B). Across the five mosquito lineages, there were 204 unique SNVs that rose slowly over the ten passages, but remained below 25% frequency. Two SNVs in lineage E that exhibited parallel trajectories indicative of genetic hitchhiking were the only SNVs to achieve greater than 50% frequency. The observed SNV trajectory patterns in mosquitoes is consistent with loose transmission bottlenecks and weak positive selection following IT inoculation, such that SNVs arise, persist, and accumulate at lower frequencies. Similar to serial mosquito passage, alternating passage did not yield any high-frequency SNVs shared across the three lineages that progressed to passage 10 (Figure 4C). One SNV, NS2B M32I, found at near 100% frequency in lineage A from passage 3 onwards, was previously associated with enhanced fetal infection in mice, but reduced transmissibility in mosquitoes (referred to as M1404I) (Lemos et al., 2020). To our knowledge, none of the other consensus-level SNVs detected in serial mosquito passage or alternating passage have been phenotyped. Unlike the accumulation of medium-frequency SNVs during serial mosquito passage, SNVs arising during alternating passage tended to either quickly rise to high frequency or be lost after a single passage. This SNV trajectory pattern in alternating passage lineages is consistent with sequential tight bottlenecks that drive SNVs to fixation or extinction.

### Differential selective pressures in mice and mosquitoes

In light of the shared SNVs arising during serial mouse passage, we sought to quantify the degree of convergent evolution in each passage type. Any variant (>1% frequency) that arose in more than one homotypic lineage was defined as convergent, and total convergence was quantified as the proportion of SNVs that were convergent (Figure 5A-B, Figure 5 — supplemental figure 1). Across all the lineages, there was a greater degree of convergent evolution during serial mouse passage (25.4%) than serial mosquito (14.8%) or alternating passage (12.5%; ***X***^2^, *P*<0.017). In the serial mosquito and alternating passage lineages, only 18.8% (6/32) and 6.5% (2/31) high-frequency SNVs (>10% frequency), respectively, were shared across multiple homotypic lineages, whereas 53.8% (7/13) were shared across multiple serial mouse lineages.

**Figure 5.**
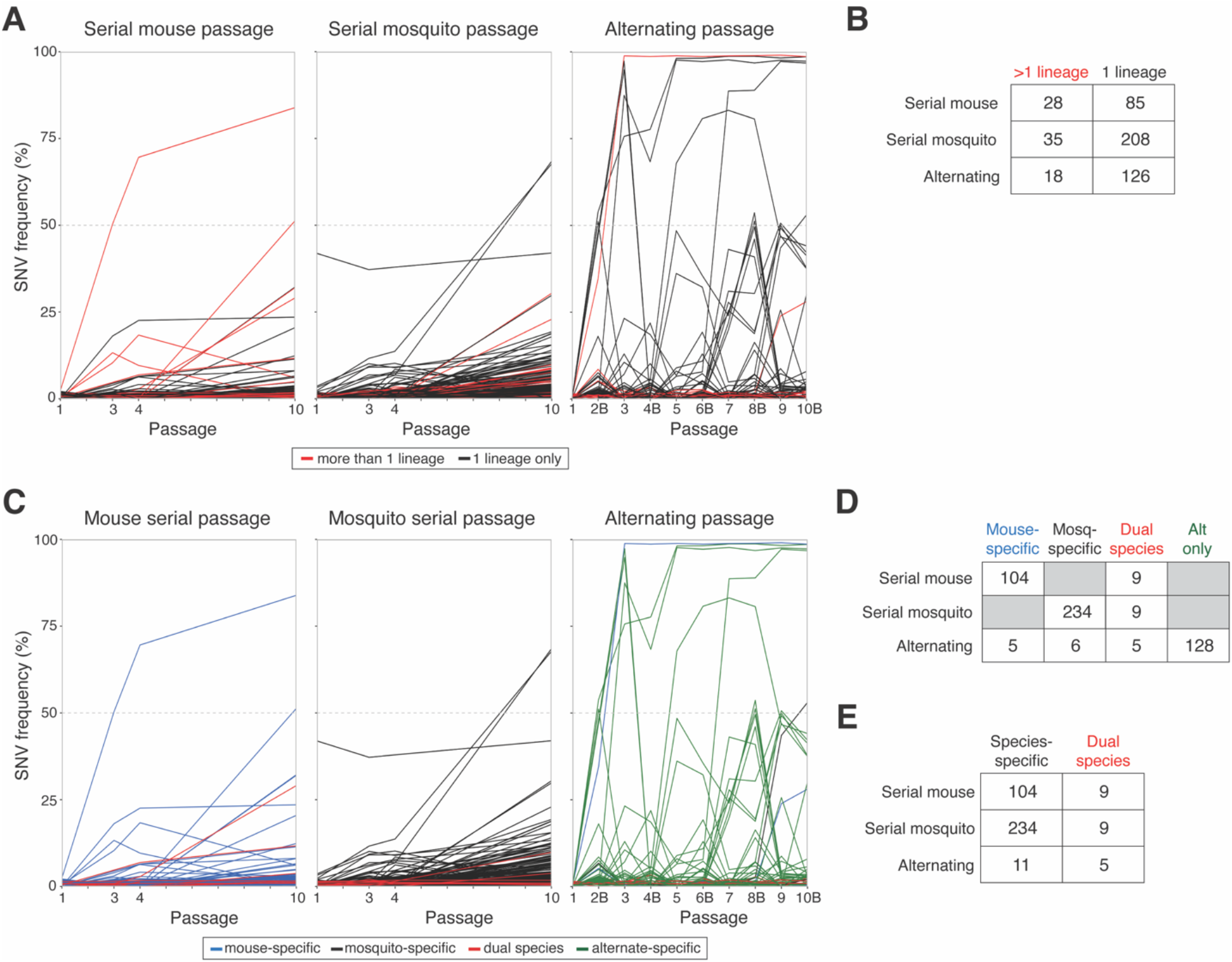
Convergence and species-specificity of individual single-nucleotide variants (SNVs) detected at >1% frequency at any passage. (**A**) Convergent SNVs, colored red, were defined as being detected in more than one homotypic lineage at any passage number. **(B)** Abundance of convergent and non-convergent SNVs for each passage series. **(C)** SNVs were defined as being mouse- or mosquito-specific if they were detected in one serial passage series and not the other. Dual species SNVs were detected in both serial mouse and serial mosquito passage series. For the alternating passage series, any SNV not defined as species-specific or dual species was defined as alternate-specific. **(D)** Abundance of species-specific, dual species, and alternate-specific SNVs in each passage series. **(E)** Abundance of species-specific and dual species SNVs in each passage series. For panels A and C, composite figures of SNV trajectories were generated by calculating the mean frequency of each SNV across all lineages it was detected in. For the alternating passage series, even passages are labeled with “B” to indicate mosquito bodies.

To understand the overlap in genetic sequence space explored by ZIKV during each passage type, we determined whether SNVs arising in the presence or absence of host-cycling were more or less likely to be species-specific. Each SNV detected during any passage at greater than 1% frequency was assigned as being mouse-specific if it only arose during serial mouse passage, and vice versa for mosquito-specific SNVs (Figure 5C, Figure 5 — supplemental figure 2). Dual-species SNVs were those detected during both serial mouse and serial mosquito passage. For alternating passage, a fourth classification, alternate-specific, was included for SNVs that only arose during alternating passage and never during serial conspecific passage. In total, more than twice as many SNVs (243 vs 113) were detected in serial mosquito passage than serial mouse passage. Despite two alternating passage lineages ending at passage 4, more SNVs were also detected during alternating passage than serial mouse passage (144 vs 113). During serial mouse passage, 92.0% of (104/113) SNVs were mouse-specific whereas only 7.9% (9/113) were dual-species (Figure 5D). Similarly, during serial mosquito passage, 96.3% (234/243) were mosquito-specific whereas only 3.7% (9/243) were dual-species (Figure 5D). There was no statistically significant difference in the high degree of SNV species specificity during serial mouse and serial mosquito passage (*X*^2^, *P*=0.09, Figure 5D). During alternating passage, the majority of SNVs were alternate-specific 88.9% (128/144) while only 3.5% (5/144), 4.2% (6/144), and 3.5% (5/144) were mouse-specific, mosquito-specific, and dual-species, respectively. Mouse-specific SNVs were similarly likely to arise during alternating passage as mosquito-specific SNVs (Fisher’s exact test, *P*=0.28; Figure 5D). In contrast, dual-species SNVs were significantly more likely to arise during alternating passage than species-specific SNVs (Fisher’s exact test, *P*<0.0001; Figure 5E). These data demonstrate very little overlap in genetic sequence space used by mouse and mosquito-adapted ZIKV, or by mouse or mosquito-adapted ZIKV and alternately passaged ZIKV. Further, the overlapping sequence space used by mouse or mosquito-adapted ZIKV and alternately passaged ZIKV was biased for mutations that arose during adaptation in both species.

In addition to trends in individual SNVs, the relative effect of selective and stochastic mechanisms were compared between passage types using population-level metrics to better understand if host environments have different impacts on virus populations. Overall genome-wide population diversity, assessed by nucleotide diversity (pi), was comparable at passages 1 through 4 across all passage series (two-way ANOVA with Tukey’s post-hoc comparisons, *P*>0.05; Figure 6A). By passage 10, serial mosquito lineages exhibited greater diversity than serial mouse lineages, which in turn were more diverse than alternating passage lineages (two-way ANOVA with Tukey’s post-hoc comparisons, *P*<0.028; Figure 6A). The mutational spectra at passage 10 did not appear biased by technical artifacts, with transition substitutions occurring at a significantly greater frequency than transversion substitutions across all passage series (matched one-way ANOVA with Dunnett’s post-hoc comparisons, *P*<0.01; Figure 6 — supplemental figure 1). The diversity at non-synonymous versus synonymous sites (piN/piS) was employed as a proxy measurement for natural selection pressures, with values greater than 1 indicative of positive selection and less than 1 indicative of purifying selection. Mean piN/piS values were consistently less than 1 for all passage types, but were significantly higher in serial mouse passages, suggesting that purifying selection was more relaxed in mice (two-way ANOVA, *P*=0.032; Figure 6B). Lastly, adaptation to host dinucleotide usage biases was compared between passage types by tracking changes in CpG and UpA dinucleotide usage within virus populations. In mice and humans, CpG and UpA dinucleotides are suppressed within the host genome (Sexton and Ebel, 2019), whereas only UpA dinucleotides are suppressed within mosquito genomes. In line with host and vector biases, CpG usage was strongly suppressed after the first serial passage in mice (linear regression non-zero slope F test, *P*=0.0060), but no suppression was observed during serial mosquito or alternating passage (linear regression slope>0; Figure 6C). Similarly in agreement with host and vector biases, UpA usage was suppressed over serial passages in both mice and mosquitoes (linear regression non-zero slope F tests, *P*<0.0001), but not over alternating passages (linear regression slope>0; Figure 6D).

**Figure 6.**
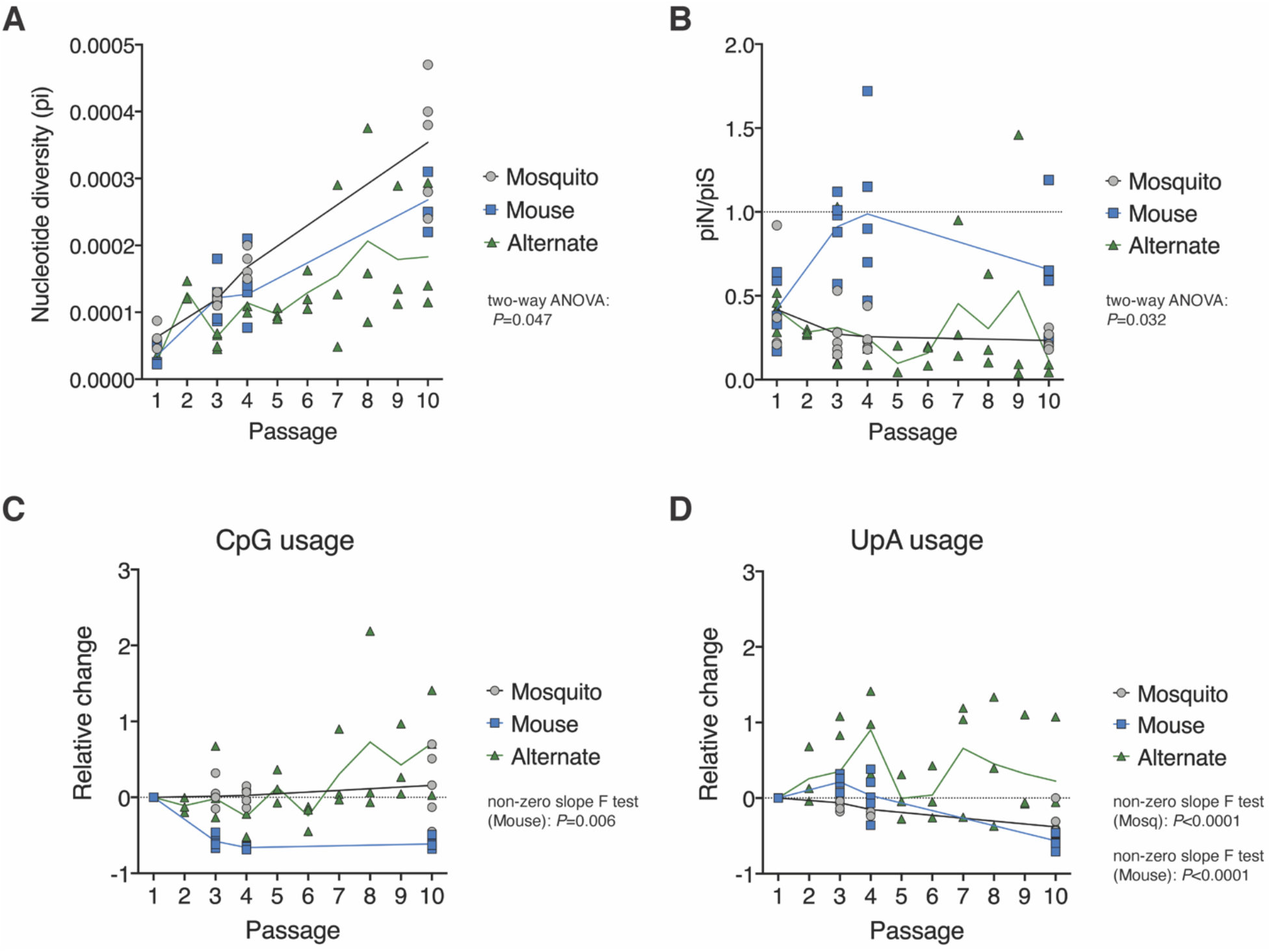
Genome-wide patterns of genetic diversity and dinucleotide usage. **(A)** Genome-wide nucleotide diversity (pi) in serial and alternate passage series over sequential passages. **(B)** Mean ratio of nucleotide diversity at nonsynonymous sites (piN) and synonymous sites (piS) across the genome. **(C)** Change in CpG usage relative to passage 1 in serial and alternate passage series. **(D)** Change in UpA usage relative to passage 1 in serial and alternate passage series. For panels A and B, two-way ANOVA *P*-values are provided for comparison of all three groups at passage 1, 3, 4, and 10. For panels C and D, only statistically significant *P*-values from linear regression non-zero slope F tests are provided. In all panels, solid lines represent mean values over sequential passages. Deep sequencing data was available for all 10 alternating passages, but only passages 1, 3, 4, and 10 of serial passage series. For serial passage series, n=5 lineages at each passage. For alternating passage, n=5 lineages at passages 1-3 and n=3 lineages for passages 4-10. Even numbered alternating passages are mosquito body samples. Alternate passage 2 data are from single mosquitoes that contributed to onward transmission, while alternate passage 4, 6, 8, and 10 are pools of all infected mosquitoes.

## Discussion

By naturally cycling between humans and mosquitoes, mosquito-borne viruses must alternately navigate distinct host environments and barriers to infection and transmission that restrict virus evolution. Unlike other mosquito-borne viruses, ZIKV frequently bypasses mosquitoes and is directly transmitted human-to-human from mother to fetus or between sexual partners. Infants infected *in utero* are likely dead-end hosts who do not contribute to onward transmission, but people infected by sexual transmission develop systemic infections and may transmit onwards to mosquitoes or additional sexual partners. Thus, direct human-to-human sexual transmission potentially enables ZIKV to redirect its evolutionary trajectory and quickly adapt to humans. In the current study, we model direct and alternating ZIKV transmission chains in mosquitoes and mice to elucidate the evolutionary pressures at play and the potential for vertebrate host-adaptation under different transmission conditions. We show that directly transmitted ZIKV rapidly adapts to mice resulting in faster and higher rates of mortality. The rise in virulence occurs in concert with the acquisition of one or two viral mutations of single amino acid, NS2A A117V or A117T. In contrast, ZIKV virulence does not increase during natural host-cycling by bloodfeeding transmission and the NS2A mutations are never detected.

In two strains of American-sublineage ZIKV not used here, NS2A A117V was previously shown to confer enhanced virulence in mice (Ávila-Pérez et al., 2019), but the phenotypic effect of NS2A A117T has yet to be determined and warrants further investigation. The consistent emergence of the NS2A 117 substitutions coincidental with the sudden loss of barcode diversity is evidence for selective sweeps, as opposed to genetic bottlenecks where population diversity is lost, but substitutions emerge randomly. Barcode sharing between serial mouse lineages after the selective sweeps indicates that the polymorphisms were likely present at below our 1% frequency cut-off in the ZIKV-BC stock and did not always arise *de novo* in mice. Interestingly, neither NS2A 117 substitution was detected in any serial mosquito or alternating passage. Two other uncharacterized substitutions—NS2A I139T and NS4A E19G— consistently arose during serial mouse passage, with only the former detected at low frequency in two serial mosquito lineages. The repeated emergence of these two substitutions, without aligned trajectories indicative of genetic hitchhiking, suggests they may confer fitness advantages, but confirmatory phenotypic analyses are warranted. These findings indicate that restriction of ZIKV host-adaptation during natural host-cycling is robust, with near complete suppression of known and presumptive beneficial mutations consistently selected for when host-cycling is circumvented. Thus, it is possible that releasing ZIKV from host-cycling through direct human-to-human transmission may reduce the barrier for emergence of virulence-enhancing mutations.

Genetic drift and strong purifying selection appeared to be the predominant evolutionary forces during serial mosquito and alternating passage. This is supported by the paucity of convergent SNVs and lower genetic diversity at nonsynonymous sites than synonymous sites across the ZIKV genome. In contrast, there is evidence for directional selection and weak purifying selection during serial mouse passage, with a greater proportion of convergent SNVs and near equal diversity at nonsynonymous and synonymous sites. These findings are in line with evolutionary forces acting on chikungunya virus (genus *Alphavirus*, family *Togaviridae*) (Riemersma et al., 2019; Riemersma and Coffey, 2019) and dengue virus (genus *Flavivirus*, family *Flaviviridae*) (Lequime et al., 2017, 2016; Vasilakis et al., 2009), but oppose trends observed with West Nile virus (genus *Flavivirus*, family *Flaviviridae*) (Deardorff et al., 2011; Grubaugh et al., 2017; Grubaugh and Ebel, 2016). The stochastic ZIKV evolution observed during alternating passage is, in large part, the result of tight genetic bottlenecks in mosquitoes during peroral infection and salivary transmission. Here, and in our previous studies (Aliota et al., 2018; Weger-Lucarelli et al., 2018), barcoded ZIKV clearly highlights these bottlenecks via sequential, drastic reductions in barcode abundance in mosquito bodies and saliva. Furthermore, individual SNV trajectories during alternating passage display strong founder biases with SNVs either rising to dominance or being lost after mosquito infection. Bottleneck effects are not evident in ZIKV evolution during serial mosquito passage due the use of IT inoculation that bypasses the bottleneck sites in the midgut and salivary glands. IT inoculation was employed to model vertical transmission in mosquitoes, and was additionally necessitated by the improbability of orally infecting naive mosquitoes with infectious, low-titer mosquito saliva.

Despite the preponderance of evidence for genetic drift and purifying selection during serial mosquito passage, we demonstrate that directional selection acts on dinucleotide usage with UpA, but not CpG, dinucleotide usage being suppressed in mosquitoes. In serial mouse passage, we observed similar suppression of UpA dinucleotides, but saw even more stringent suppression of CpG dinucleotides. The observed ZIKV dinucleotide usage patterns in mice and mosquitoes align with the dinucleotide usage biases in each host (Lobo et al., 2009; Sexton and Ebel, 2019). Mosquitoes and vertebrates exhibit disparate dinucleotide usage biases in their transcriptomes, and arbovirus genomes typically adopt an intermediate usage pattern at consensus-level compared to single-host viruses, presumably to accommodate both vector and vertebrate host environments (Halbach et al., 2017; Sexton and Ebel, 2019). To our knowledge, this is the first evidence of the dinucleotide selective pressure acting on a multi-host virus at the sub-consensus level, as opposed to the consensus level. Unsurprisingly, we observed no clear trends in dinucleotide usage patterns during natural alternating passage, almost certainly the byproduct of bottleneck events. This provides further evidence that natural alternating transmission restricts ZIKV’s capacity to adapt to vertebrate hosts and also mosquito vectors.

The fitness trade-off hypothesis posits that fitness gains in one host come at the cost of fitness losses in the other host. Here, we demonstrate evidence to the contrary. Serial passage of ZIKV in mosquitoes increased viral replication in mosquitoes, but replication was not reduced in mice. These data are more consistent with the notion that the degree of host specialization can dramatically alter the evolution of virulence in pathogen populations, and that a fitness gain in one environment may paradoxically broaden the overall phenotypic potential of a virus. To this end we are performing additional studies to evaluate the phenotypic effects of host-cycling release in mosquitoes to determine if the lack of fitness trade-offs are observed in both hosts. Antagonistic pleiotropy—contradictory phenotypes for the same mutations in hosts and vectors—is the rationale underlying the fitness trade-off hypothesis. Interestingly, although our data refute fitness trade-offs for host-adapted lineages, there is evidence for antagonistic pleiotropy in that very few SNVs arise during both mouse and mosquito-adaptation, and very few species-specific SNVs arise during alternating transmission. This suggests that antagonistic pleiotropy can exist without apparent fitness trade-offs for the ZIKV population, possibly due to weak antagonism and overlapping viable fitness landscapes (Coffey et al., 2013; Novella et al., 2011). That said, species-specific SNVs (potentially antagonistic) were less likely to arise in alternately passaged ZIKV than dual species SNVs that presumably had neutral or beneficial fitness effects in both hosts. Overall, alternately passaged ZIKV exhibited very little overlap in SNV usage with the mouse- or mosquito-adapted ZIKV. Non-mutually exclusive explanations for the uniqueness of alternately passaged ZIKV are: 1) genetic drift in broad, viable sequence space such that few mutations are shared by chance, and 2) that host-cycling acts as a selection pressure pushing the ZIKV population into a unique region of sequence space. Further investigations into the relative contribution of each explanation are worthwhile because the likelihood of host-adaptive SNVs emerging is likely higher in the first scenario than the second. Additionally, making that distinction would inform whether ZIKV’s evolutionary potential under natural host-cycling conditions can be explored or predicted by experimental adaptation to animal models or mosquitoes.

Taken together, our findings clarify the effect of host-cycling on ZIKV evolution and highlight the potential for rapid host-adaptation with direct vertebrate transmission chains. Whether similar adaptation would be observed with direct human-to-human transmission remains unclear. A potential limitation of this study is that direct transmission by needle inoculation may imperfectly model sexual transmission dynamics. Further studies are therefore needed to assess ZIKV host-adaptation using animal models of sexual transmission. Nonetheless, our data suggest that prevention of direct human transmission chains should be a public health priority to thwart the emergence of virulence-enhancing mutations.

## Materials and Methods

### Cells and virus

African Green Monkey cells (Vero; ATCC CCL-81) and human embryonic kidney cells (HEK293T; ATCC CRL-3216) were cultured in Dulbecco’s modified Eagle media (DMEM; Gibco) supplemented with 10% fetal bovine serum (FBS; Cytiva HyClone), 2 mM L-glutamine, 1.5 g/L sodium bicarbonate, 100 U/ml penicillin, and 100 μg/ml of streptomycin at 37C in 5% CO_2_. Larval *Aedes albopictus* cells (C6/36; ATCC CRL-1660) were cultured in DMEM supplemented with 10% FBS, 2 mM L-glutamine, 1.5 g/L sodium bicarbonate, 100 U/ml penicillin, and 100 μg/ml of streptomycin at 28°C in 5% CO_2_. The barcoded ZIKV infectious clone was constructed by bacteria-free cloning of the ZIKV PRVABC59 strain genome (GenBank: KU501215.1), as previously described (Aliota et al., 2018; Weger-Lucarelli et al., 2018). Briefly, the ZIKV PRVABC59 isolate was passaged three times on Vero cells and twice on C6/36 cells, followed by PCR amplification of the whole genome in two overlapping amplicons. The genetic barcode, degenerate nucleotides at the third position of 8 consecutive codons in *NS2A* (Figure 1A), was then introduced via an overlapping PCR-amplified oligo. The two amplicons were assembled with a 5’ CMV promoter amplified from pcDNA3.1 (Invitrogen) by Gibson assembly (New England Biosciences (NEB)) followed by enzymatic digestion of remaining ssDNA and non-circular dsDNA. Full-length ZIKV constructs were amplified using rolling circle amplification (Qiagen repli-g mini kit) and genomic integrity verified by restriction digestion and Sanger sequencing. Infectious barcoded ZIKV (ZIKV-BC) rescue was performed in HEK293T cells.

### Virus titration

Infectious virus was titrated by plaque assay on Vero cells. A confluent monolayer of Vero cells were inoculated with a 10-fold dilution series of each sample in duplicate. Inoculated cells were incubated for 1 hour at 37°C and then overlaid with a 1:1 mixture of 1.2% oxoid agar and 2X DMEM (Gibco) with 10% (vol/vol) FBS and 2% (vol/vol) penicillin/streptomycin. After four days, the cell monolayers were stained with 0.33% neutral red (Gibco). Cells were incubated overnight at 37°C and plaques were counted. Plaque counts were averaged across the two replicates and the concentration of infectious ZIKV was back-calculated from the mean.

Viral RNA was isolated directly from mouse serum, mosquito saliva collected in cell culture media, and cell culture supernatant. Mosquito bodies were homogenized in PBS supplemented with 20% FBS and 2% penicillin/streptomycin with 5mm stainless steel beads with a TissueLyser (Qiagen) prior to RNA isolation. RNA was isolated with the Maxwell RSC Viral Total Nucleic Acid Purification Kit on a Maxwell RSC 48 instrument (Promega). Isolated ZIKV RNA was titrated by qRT-PCR using TaqMan Fast Virus 1-Step Master Mix (ThermoFisher) and a LightCycler 480 or LC96 instrument (Roche). Final reaction mixtures contained 600 nM of each ZIKV-specific qRT-PCR primer (5’-CGY TGC CCA ACA CAA GG-3’ and 5’-CCA CYA AYG TTC TTT TGC ABA CAT-3’) and 100 nM of probe (5’-6-FAM-AGC CTA CCT TGA YAA GCA RTC AGA CAC YCA A-BHQ1-3’) (Lanciotti et al., 2008). Cycling conditions were as follows: 50°C for 5 minutes, 95°C for 20 seconds, and 50 cycles of 95°C for 15 seconds followed by 60°C for 1 min. ZIKV RNA titers were interpolated from a standard curve of diluted *in vitro* transcribed ZIKV RNA. The limit of detection for this assay is 100 ZIKV genome copies/ml.

### Mice and mosquitoes

*Ifnar1*^−/−^ mice on the C57BL/6 background were bred in the specific pathogen-free animal facilities of the University of Wisconsin-Madison (UW) Mouse Breeding Core within the School of Medicine and Public Health, or in the specific pathogen-free animal facilities of the University of Minnesota (UMN) College of Veterinary Medicine. Three to six week-old mice of mixed sex were used for all experiments.

*Ae. aegypti* mosquitoes used in this study were maintained at UW and UMN using previously described rearing protocols (Christensen and Sutherland, 1984). The *Ae. aegypti* line used in this study was established from several hundred eggs collected from ovitraps placed around the municipality of Buenos Aires (communa no. 9), a southeast suburb of Medellin, Colombia. Mosquitoes used in this study were from generations 3 to 30 of the laboratory colony. Three-to six-day-old female mosquitoes were used for all experiments.

This study was approved by the UW and UMN Institutional Animal Care and Use Committees (Animal Care and Use Protocol Numbers V5519 (UW) and 1804–35828 (UMN)).

### Serial passage in mice or mosquitoes

Five *Ifnar1*^−/−^ mice were subcutaneously inoculated in the left hind footpad with 10^3^ PFU of ZIKV-BC stock as passage 1 of five replicate lineages. Submandibular blood draws were performed two days post-inoculation (dpi) and serum was processed for virus titration, sequencing, and for onward passaging. Serial passaging for each lineage was maintained by inoculating a naive mouse with the 2 dpi serum diluted to 10^3^ PFU for ten total passages.

Female *Ae. aegypti* mosquitoes were anesthetized on ice and intrathoracically inoculated (Hong et al., 2003) with 100 PFU of ZIKV-BC in 1 ul. Inoculated mosquitoes were maintained on 0.3 M sucrose in an environmental chamber at 26.5°C ± 1°C, 75% ± 5% relative humidity and with a 12-hour photoperiod within the Department of Pathobiological Sciences Biosafety Level 3 insectary at UW. At 12 dpi, mosquitoes were individually homogenized in 1ml of PBS supplemented with 20% FBS and 2% penicillin/streptomycin. The supernatant was then collected and used for virus titration, sequencing, and onward passaging. Supernatant from five individual mosquitoes was then used to serially passage 100 PFU of virus through five replicate lineages of mosquitoes for ten passages.

### Alternating passage

Five *Ifnar1*^−/−^ mice were subcutaneously inoculated in the left hind footpad with 10^3^ PFU of ZIKV-BC. Two days post inoculation, mice under ketamine/xylazine anesthesia were fed on by cartons of female *Ae. aegypti* that had been sucrose starved for 14-16 hours prior to mouse feeding. After mosquito bloodfeeding, submandibular blood draws were performed to collect serum for virus titration and sequencing. Mosquitoes were anesthetized on ice and mosquitoes that fed to repletion were selected and placed in new cartons containing an oviposition cup. Bloodfed mosquitoes were maintained on 0.3 M sucrose in an environmental chamber at 26.5°C ± 1°C, 75% ± 5% relative humidity and with a 12-hour photoperiod within the Veterinary Isolation Facility BSL3 Insectary at UMN. 12 days-post-feeding and following oviposition between 8-10 days, mosquitoes bloodfed on new *Ifnar1*^−/−^ mice. Mosquitoes were then triethylamine anesthetized and saliva and whole bodies were collected from those that fed to repletion for sequencing and virus titration. Two days post bloodfeeding, these mice were fed on by naive cartons of mosquitoes to continue alternate passaging through ten passages.

### Library preparation and sequencing

Virus barcode libraries for the mouse and mosquito serial passage samples were generated with unique molecular identifiers (UMI) to filter sequencer and PCR errors that could produce false barcode sequences. ZIKV RNA concentrations in the alternating passage samples were too low to employ the UMI approach, so viral barcodes were sequenced by the whole genome sequencing (WGS) approach described below. UMIs consisted of 12 random nucleotides inserted into the reverse primer used for reverse transcription of the ZIKV barcode region (5’-GGA GTT CAG ACG TGT GCT CTT CCG ATC TNN NNN NNN NNN NCC CCC GCA AGT AGC AAG GCC TG-3’). UMI-tagged cDNA was treated with RNase H, purified with magnetic beads (Agencourt RNAclean XP), and then PCR amplified for 20 cycles (NEB Phusion Master Mix; Forward: 5’-TCT TTC CCT ACA CGA CGC TCT TCC GAT CTT GGT TGG CAA TAC GAG CGA TGG TT-3’, Reverse: 5’-GTG ACT GGA GTT CAG ACG TGT GCT CTT CC-3’). Amplicons were bead-purified (Agencourt Ampure XP) and then PCR amplified for 34 additional cycles using the same reverse primer and a forward primer bearing a 6-nucleotide index sequence (Forward: 5’-CAA GCA GAA GAC GGC ATA CGA GAT NNN NNN GTG ACT GGA GTT CAG ACG TGT GCT CTT-3’). Reconditioning PCR using 1/10th volume of the unpurified index amplicon was performed for 3 cycles using the same reagents as the index PCR reaction. The entire volume of reconditioned UMI barcode libraries were purified by gel extraction (Qiagen QIAquick Gel Extraction Kit).

Whole genome ZIKV sequencing libraries were generated with a previously described tiled PCR amplicon approach (Grubaugh et al., 2019; Quick et al., 2017). Briefly, 10^6.15^ ZIKV genome copies were converted to cDNA with Superscript IV reverse transcriptase and random hexamer primers (ThermoFisher). PCR amplification of the entire ZIKV coding region was then performed in two reactions with pools of non-overlapping PCR primer sets. Technical duplicates were generated for each WGS and UMI barcode library. All libraries were quantified by a Qubit 3 fluorometer (ThermoFisher) and quality assessed by Agilent Bioanalyzer prior to sequencing. UMI barcode libraries were sequenced with paired-end 250 base pair reads on an Illumina MiSeq (Illumina MiSeq Reagent Kit v2). WGS libraries were sequenced with paired-end 150 base pair reads on an Illumina NovaSeq 6000 by the UW Biotechnology Center (Illumina NovaSeq 6000 S1 Reagent Kit v1.5).

### Bioinformatic analyses

For UMI barcode sequence data, a pipeline was generated to process raw Illumina reads, extract consensus UMI reads, and calculate unique barcode abundance and frequencies. Briefly, raw paired-end reads were adapter- and quality-trimmed (q35), merged, cropped, then quality-filtered based on average base quality. High-quality reads were then grouped by UMI sequences and, for UMI groups with at least 3 reads, the consensus sequence was extracted. The 24-nucleotide barcode sequence was then extracted from all consensus sequences without ambiguous bases. Finally, the abundance and frequency of each unique barcode sequence was calculated. For mosquito samples in alternating passage lineages, concentrations of ZIKV RNA were too low to use the UMI barcode library approach, so instead, WGS data were used. First, reads were adapter- and quality-trimmed, then any paired-end reads with mismatched bases in their overlapping sequences were filtered prior to merging. High-quality merged reads were aligned to the ZIKV PRVABC59 reference sequence and reads fully covering the barcode region were isolated. Barcode sequences were then extracted from the reads, and the abundance and frequency of unique barcodes was calculated. For both the UMI and WGS barcode approach, mean barcode abundance and frequency was calculated across technical duplicate libraries and used in subsequent analyses.

For WGS data, a pipeline was generated to process raw Illumina reads, align reads at a normalized depth, call variants, and calculate diversity and dinucleotide usage metrics. Briefly, raw paired-end reads were adapter-trimmed, then any paired-end reads with mismatched bases or less than a 50-base pair overlap were filtered prior to merging. Next, merged reads were quality-trimmed and primer sequences from the tiled primer sets were trimmed from the ends of the high-quality merged reads. Reads were then aligned to the ZIKV PRVABC59 reference and normalized to a coverage depth of approximately 2500. Consensus sequences were extracted and variants were called against both the reference and consensus sequences with LoFreq* (Wilm et al., 2012). As with barcodes frequencies, variant frequencies were averaged across technical duplicate libraries and mean frequencies were used for data analyses. Genome-wide and site-specific nucleotide diversities (pi, piN, piS) were calculated with SNPGenie (v3, minfreq=0.003) (Nelson et al., 2015). Dinucleotide usage was calculated as the net change in frequency of each dinucleotide with a bespoke R script. First, dinucleotide sites were defined for each nucleotide pair across the reference genome, then potential dinucleotide sites were identified as nucleotide pairs that differ from the target dinucleotide by one nucleotide (for example, CpC or ApG for CpG dinucleotides). Then, per site dinucleotide losses were calculated as the mean frequency of point mutations at each dinucleotide site, and per site dinucleotide gains were calculated as the mean frequency of point mutations that generated the target dinucleotide at potential dinucleotide sites. Net dinucleotide usage for each dinucleotide was calculated as the per site dinucleotide gains divided by per site dinucleotide losses.

All bespoke data processing, analysis, and visualization scripts are publicly available on GitHub (https://github.com/tcflab/ZIKVBC_HostCycling). Read quality-trimming and cropping were conducted with Trimmomatic (v0.39) (Bolger et al., 2014), cutadapt (v2.3) (Martin, 2011), and fastp (v0.20.0) (Chen et al., 2018). Read merging, alignment normalization, and barcode counting were performed with BBTools (v34.48; Joint Genome Institute). Reference alignment of reads was completed with the Burrows-Wheeler Aligner (bwa-mem; v0.7.16) (Li and Durbin, 2009). Parameter settings for each process not included in the text are provided in the aforementioned scripts.

### Statistical analyses

All statistical analyses were conducted using GraphPad Prism 8 (GraphPad Software, CA, USA). Statistical significance was designated to *P* values of less than 0.05.

## Data availability

Raw Illumina sequencing data are available on the NCBI Sequence Read Archive under Bioproject PRJNA671510.

## Acknowledgements

The authors acknowledge the University of Minnesota, Twin Cities BSL3 Program for facilities and Neal Heuss for support. We thank Natalie Benett for her contribution in mosquito maintenance, Matthew Semler for his technical support, and Andrea Weiler and Mason Bliss for their contributions in virus titration. We also thank Jody Peter for maintenance of the *Ifnar1*^−/−^ colony at the University of Wisconsin-Madison.

---

**Figure 2 — Supplemental Figure 1.**
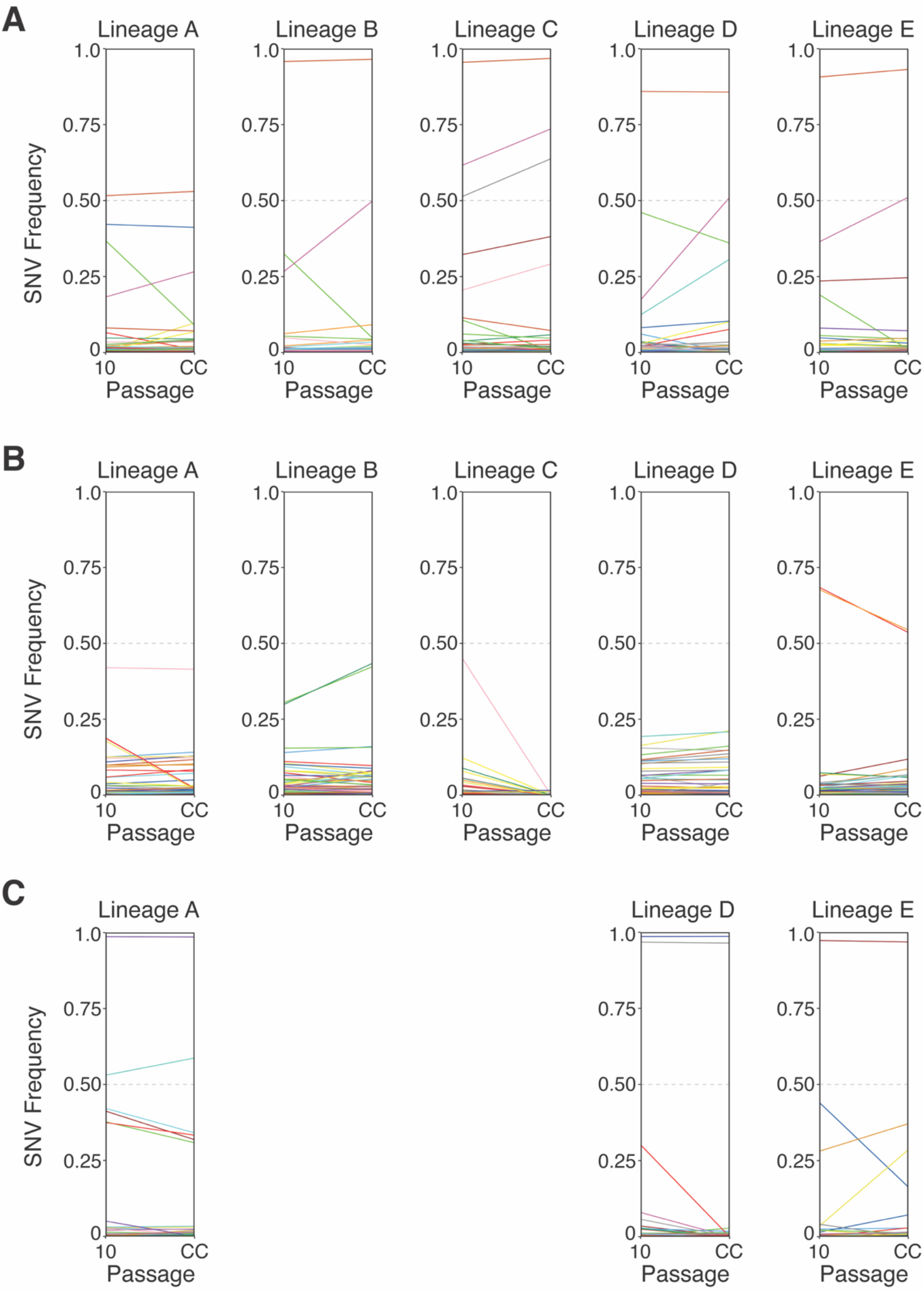
Frequencies of Zika virus single-nucleotide variants (SNVs) at passage 10 and after a single amplifying passage in cell culture. Changes in individual SNV frequencies are represented by colored lines. Colors represent the same SNVs across homotypic lineages, but are used more than once due to the large number of SNVs. SNV frequencies (0.0 to 1.0) are provided for **(A)** serial mouse passage lineages, **(B)** serial mosquito passage lineages, and **(C)** alternating passage lineages. Alternating passage lineages B and C were not maintained to passage 10, and therefore are not shown.

**Figure 3 — Supplemental Figure 1.**
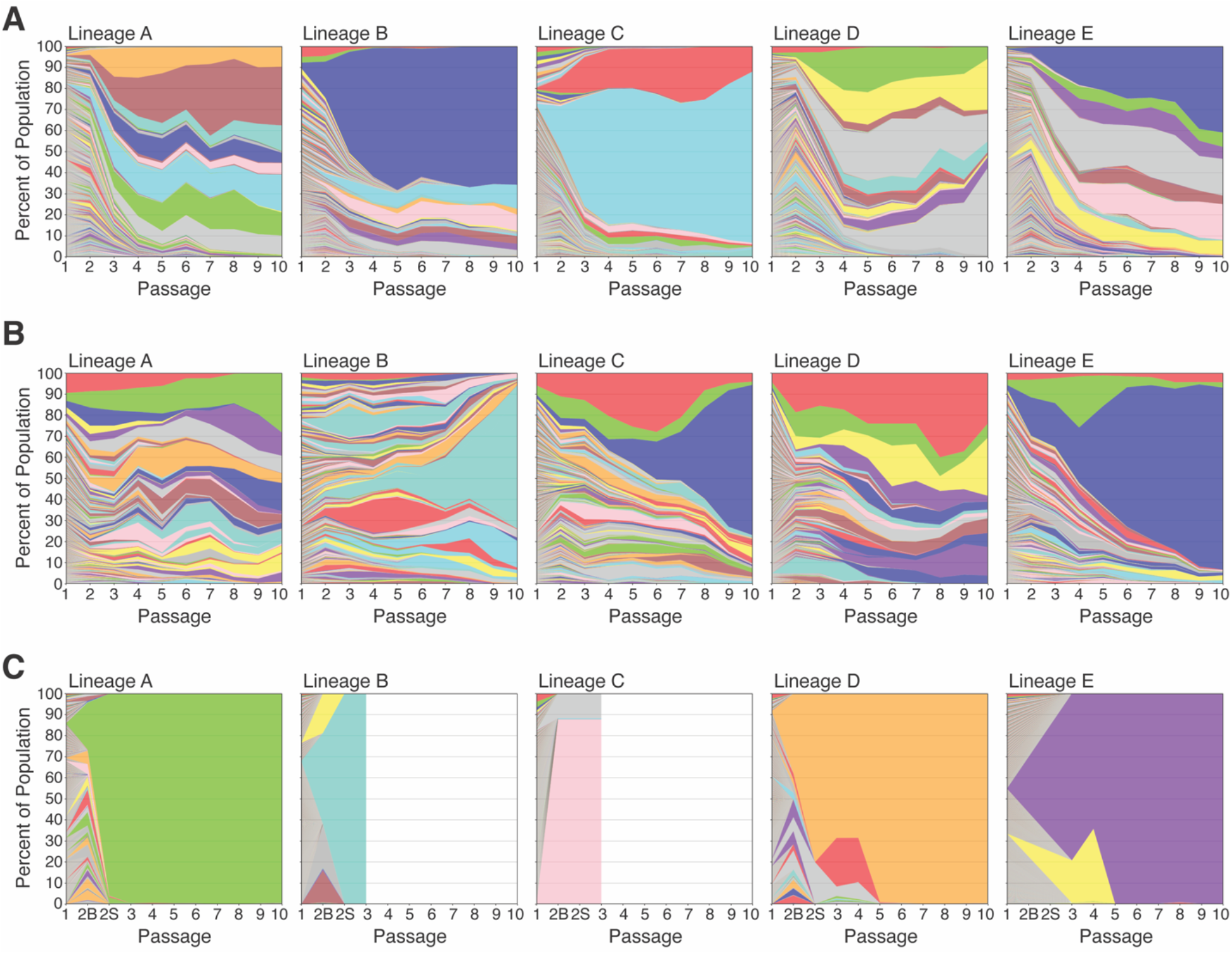
ZIKV barcode dynamics over sequential passages for individual lineages. The frequency of each barcode over serial passages are presented for. **(A)** serial mouse passage lineages, **(B)** serial mosquito passage lineages, and **(C)** alternating passage lineages. Colors represent barcode rank and are not associated with the barcode sequence. For alternating passage lineages, passage 2B and 2S are the mosquito body and saliva, respectively, that contributed to onward transmission. Alternating passage lineages B and C were not maintained past passage 3.

**Figure 3 — Supplemental Figure 2.**
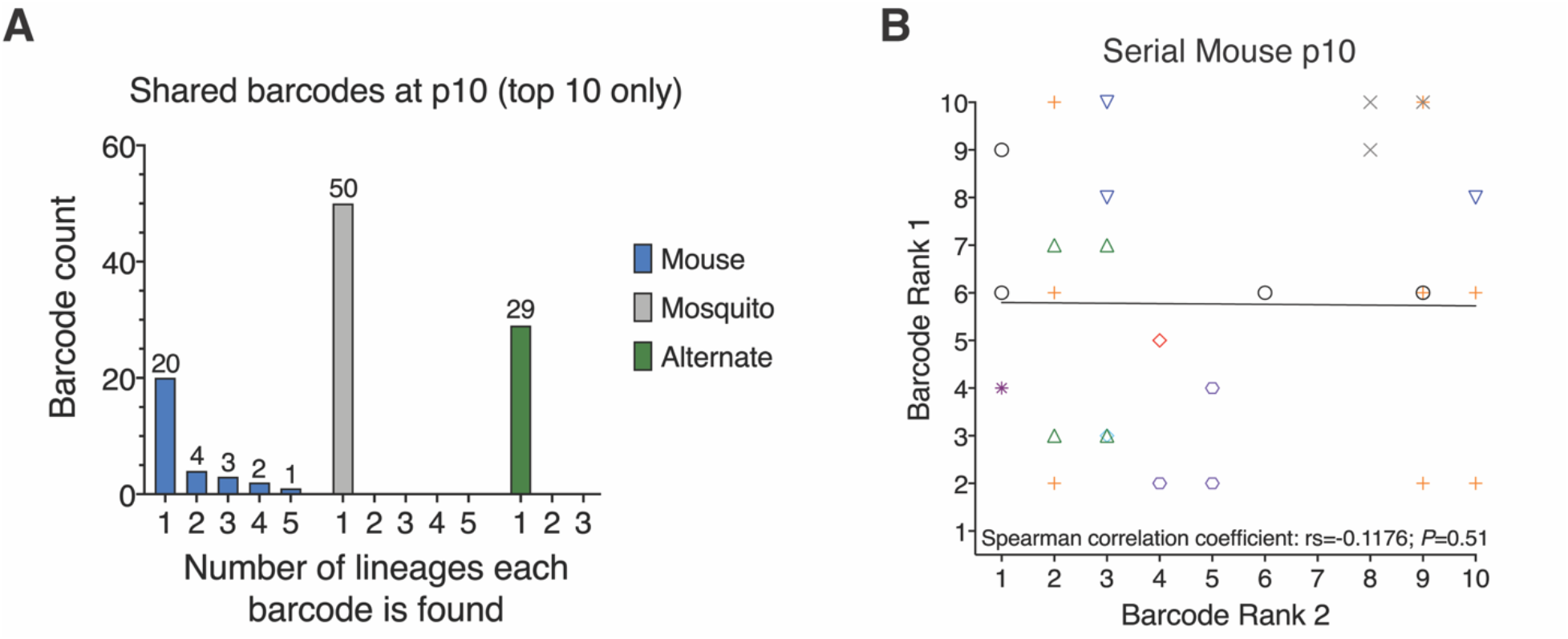
Barcodes shared between homotypic lineages at passage 10. **(A)** Histograms of the number of barcodes found in multiple (>1) lineage. Only the top 10 most frequent barcodes from each of the five lineages were analyzed (max 50 barcodes). No barcodes were shared by multiple serial mosquito or alternating passage lineages, while ten barcodes were shared by at least two serial mouse lineages. **(B)** Pairwise ranks for each of the ten barcodes shared by more than one serial mouse lineage at passage 10. Colored symbols represent the ten specific barcode sequences. The solid black line represents the line of best-fit for all pairwise ranks. The nonparametric Spearman’s correlation coefficient (rs) and *P*-value for all pairwise ranks are provided.

**Figure 4 — Supplemental Figure 1.**
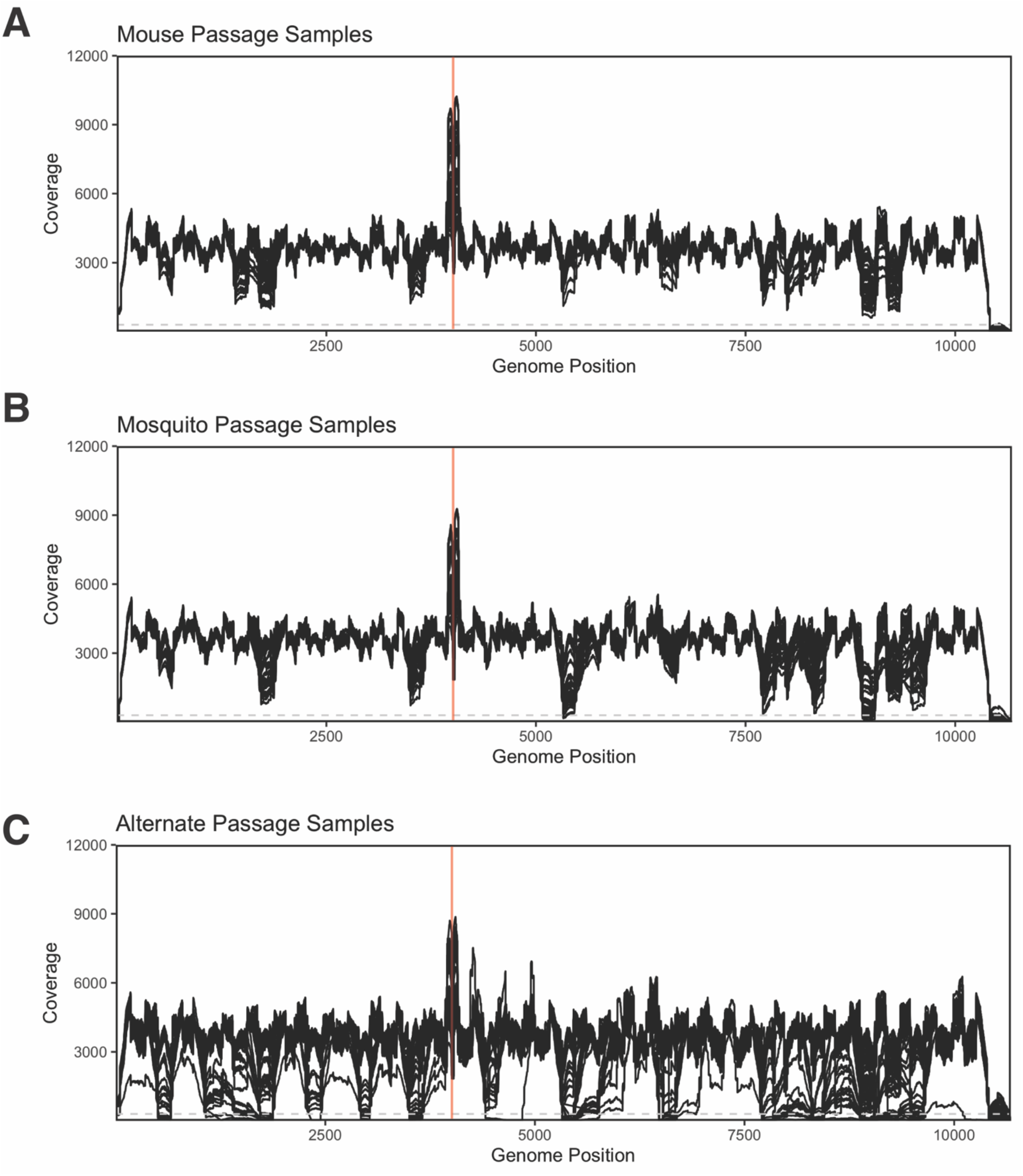
Whole-genome sequencing depth of coverage across Zika virus genome after normalization for. **(A)** serial mouse passage samples, **(B)** serial mosquito passage samples, and **(C)** alternating passage samples. All samples and replicates are individually represented by black lines. Lower coverage samples in **(C)** are predominantly from mosquito saliva. Vertical red lines indicate the location of the molecular barcode within *NS2A*. Dashed grey line at a depth of coverage of 300 reads delineates the coverage threshold used for variant calling and diversity analyses.

**Figure 5 — Supplemental Figure 1.**
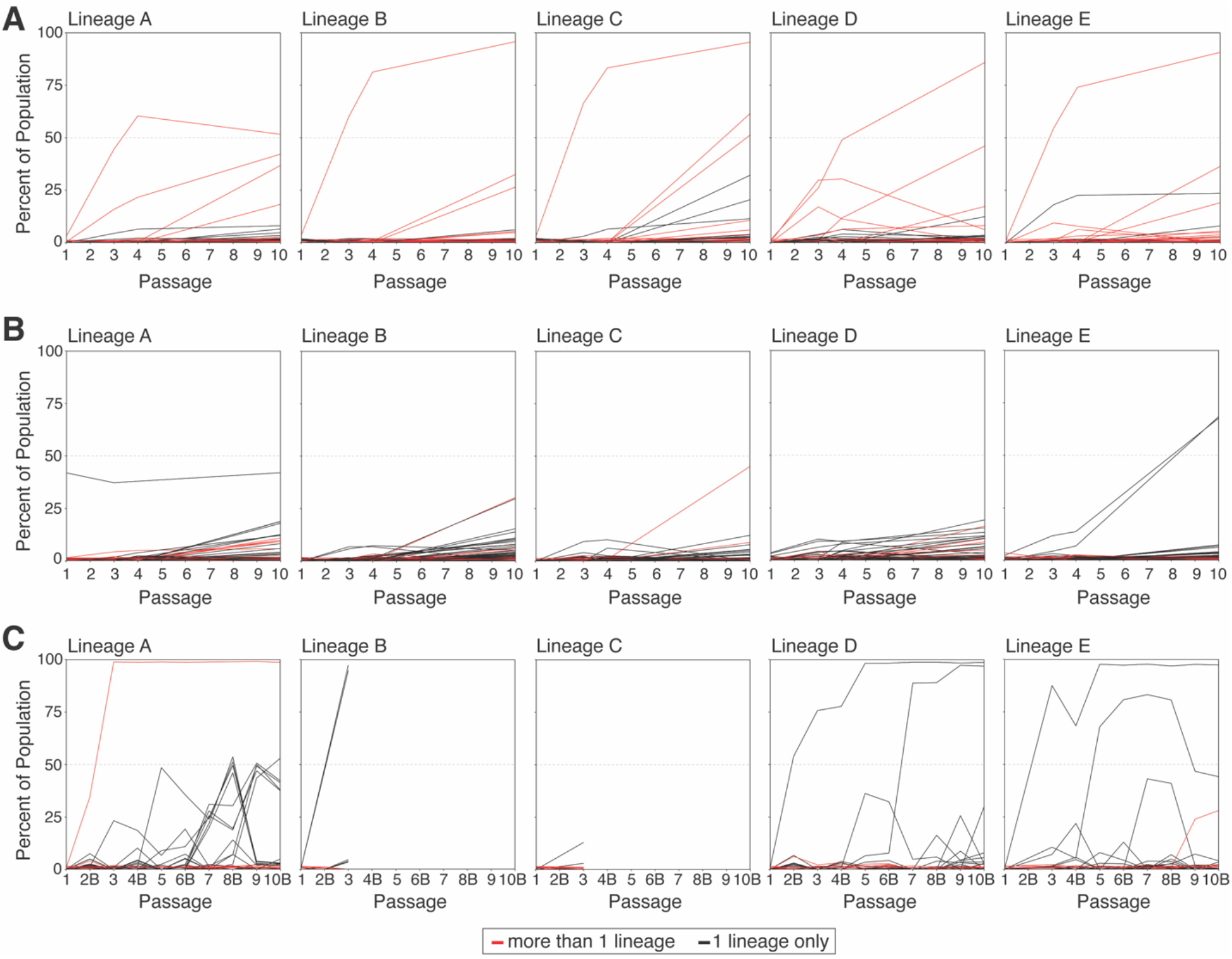
Dynamics of Zika virus single-nucleotide variants detected in more than one homotypic lineage (red) or only one lineage (black). Frequency trajectories are provided for each of the five replicate lineages of **(A)** serial mouse passage, **(B)** serial mosquito passage, and **(C)** alternating passage. Alternating passage lineages B and C were not maintained past passage 3. For alternating passage lineages in panel C, even numbered passages are noted with a “B” to indicate mosquito bodies. Alternate passage 2B is the single mosquito that contributed to onward transmission, whereas latter even-numbered passages are pools of infected mosquito bodies.

**Figure 5 — Supplemental Figure 2.**
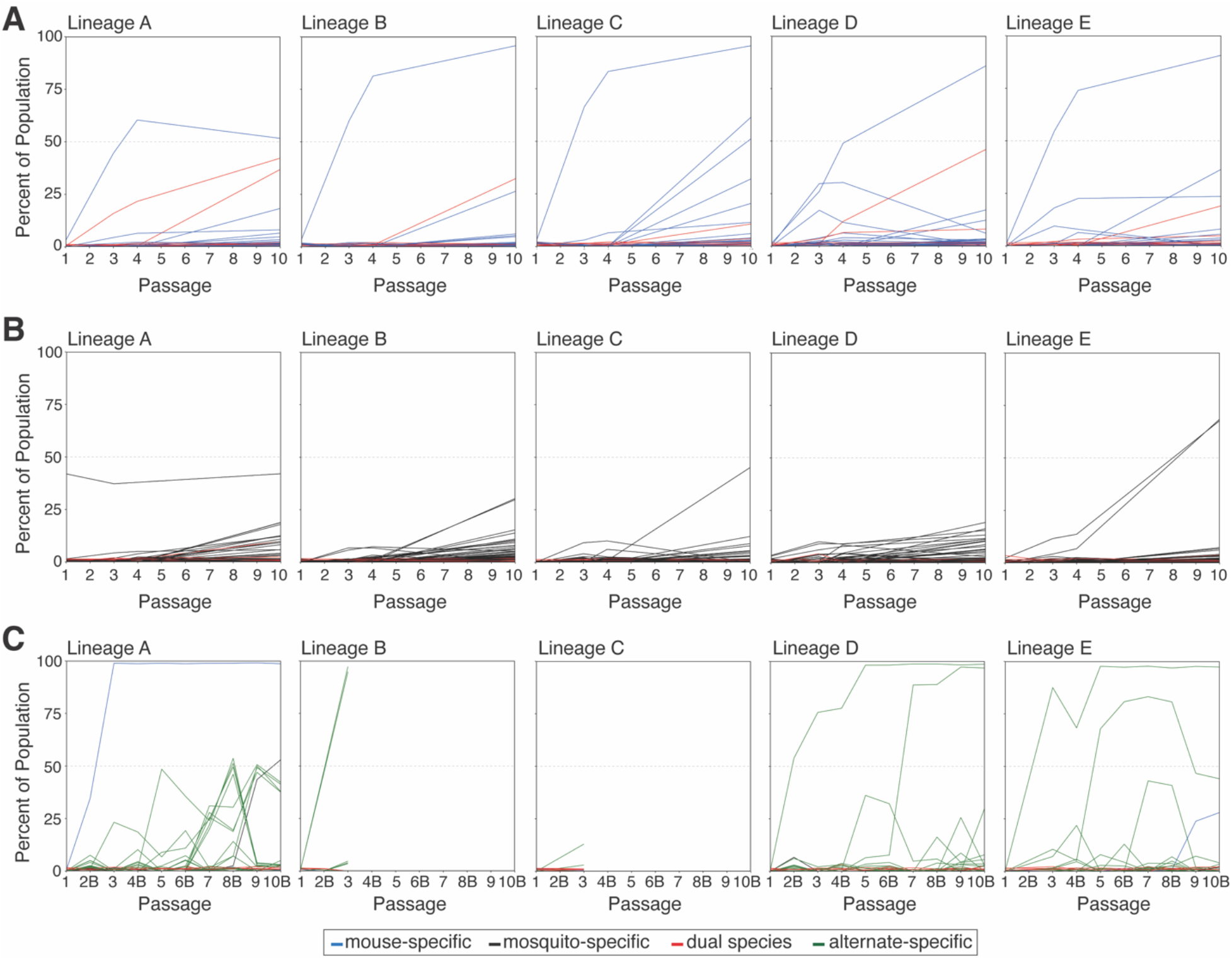
Dynamics of Zika virus single-nucleotide variants detected only in serial mouse passages (blue, “mouse-specific”), only in serial mosquito passages (black, “mosquito-specific”), in both serial mouse and serial mosquito passages (red, “dual species”), or only in alternating passages (green, “alternate-specific”). Frequency trajectories are provided for each of the five replicate lineages of **(A)** serial mouse passage, **(B)** serial mosquito passage, and **(C)** alternating passage. Alternating passage lineages B and C were not maintained past passage 3. For alternating passage lineages in panel C, even numbered passages are noted with a “B” to indicate mosquito bodies. Alternate passage 2B is the single mosquito that contributed to onward transmission, whereas latter even-numbered passages are pools of infected mosquito bodies.

**Figure 6 — Supplemental Figure 1.**
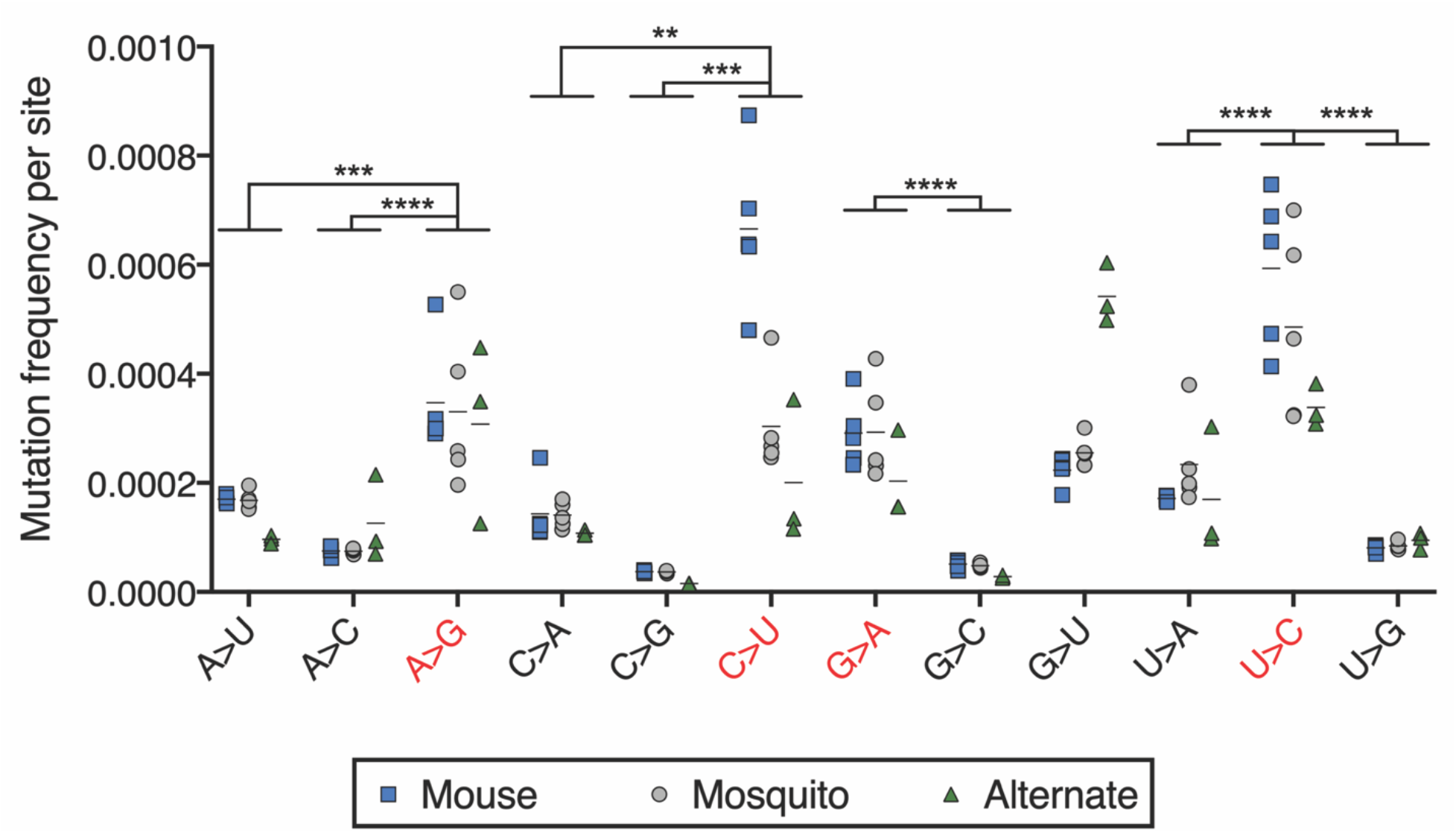
Frequency of specific nucleotide substitutions detected in all processed and aligned reads at passage 10. Frequency is reported as the average number of each substitution per site. Substitutions are shown as the reference nucleotide followed by “>” and the substitute nucleotide. Transition substitutions are highlighted with red font, whereas transversion substitutions are shown in black font. Horizontal lines represent the arithmetic means for each group. Statistically significant differences in mean frequencies of transitions versus transversions were assessed by matched one-way ANOVA tests. *P*-values less than 0.01, 0.001, and 0.0001 are denoted by **, ***, and ****, respectively. Absence of asterisks indicates a lack of statistically significant differences in mean frequencies (*P*>0.05).

**Supplemental Table 1.** Run and sample data for whole-genome sequencing. **(A)** Run metrics for whole-genome sequencing runs performed on Illumina NovaSeq. **(B)** Whole-genome sequencing library identifiers and descriptions. Q30: quality score of 30.

| Run | System | Flowcell | Clusters<br>(Raw) | Clusters<br>(PF) | % bases<br>≥Q30 | Mean base<br>quality score |
| --- | --- | --- | --- | --- | --- | --- |
| 1 | Illumina NovaSeq | 2X150 S1 | 1,615,376,266 | 807,688,133 | 95.51 | 36.26 |
| 2 | Illumina NovaSeq | 2X150 S1 | 1,747,936,596 | 873,968,298 | 93.53 | 35.89 |

| Library ID | Passage series | Passage number | Lineage | Library ID | Passage series | Passage number | Lineage |
| --- | --- | --- | --- | --- | --- | --- | --- |
| SCP1_1 | Serial mouse | 1 | A | AP1_1 | Alternating | 1 | A |
| SCP1_2 | Serial mouse | 1 | B | AP1_2 | Alternating | 1 | B |
| SCP1_3 | Serial mouse | 1 | C | AP1_3 | Alternating | 1 | C |
| SCP1_4 | Serial mouse | 1 | D | AP1_4 | Alternating | 1 | D |
| SCP1_5 | Serial mouse | 1 | E | AP1_5 | Alternating | 1 | E |
| SCP3_1 | Serial mouse | 3 | A | AP2_1_3B | Alternating | 2 | A |
| SCP3_2 | Serial mouse | 3 | B | AP2_1_3S | Alternating | 2 | A |
| SCP3_3 | Serial mouse | 3 | C | AP2_2_3B | Alternating | 2 | B |
| SCP3_4 | Serial mouse | 3 | D | AP2_2_3B | Alternating | 2 | B |
| SCP3_5 | Serial mouse | 3 | E | AP2_4_6B | Alternating | 2 | D |
| SCP4_1 | Serial mouse | 4 | A | AP2_4_6S | Alternating | 2 | D |
| SCP4_2 | Serial mouse | 4 | B | AP3_1 | Alternating | 3 | A |
| SCP4_3 | Serial mouse | 4 | C | AP3_2 | Alternating | 3 | B |
| SCP4_4 | Serial mouse | 4 | D | AP3_3 | Alternating | 3 | C |
| SCP4_5 | Serial mouse | 4 | E | AP3_4 | Alternating | 3 | D |
| SCP10_1 | Serial mouse | 10 | A | AP3_5 | Alternating | 3 | E |
| SCP10_2 | Serial mouse | 10 | B | AP4_1B | Alternating | 4 | A |
| SCP10_3 | Serial mouse | 10 | C | AP4_4B | Alternating | 4 | D |
| SCP10_4 | Serial mouse | 10 | D | AP4_5B | Alternating | 4 | E |
| SCP10_5 | Serial mouse | 10 | E | AP5_1 | Alternating | 5 | A |
| SCP10_A | Serial mouse | CC | A | AP5_4 | Alternating | 5 | D |
| SCP10_B | Serial mouse | CC | B | AP5_5 | Alternating | 5 | E |
| SCP10_C | Serial mouse | CC | C | AP6_1B | Alternating | 6 | A |
| SCP10_D | Serial mouse | CC | D | AP6_4B | Alternating | 6 | D |
| SCP10_E | Serial mouse | CC | E | AP6_5B | Alternating | 6 | E |
| MP1_1 | Serial mosquito | 1 | A | AP7_1 | Alternating | 7 | A |
| MP1_2 | Serial mosquito | 1 | B | AP7_4 | Alternating | 7 | D |
| MP1_3 | Serial mosquito | 1 | C | AP7_5 | Alternating | 7 | E |
| MP1_4 | Serial mosquito | 1 | D | AP8_1B | Alternating | 8 | A |
| MP1_5 | Serial mosquito | 1 | E | AP8_4B | Alternating | 8 | D |
| MP3_1 | Serial mosquito | 3 | A | AP8_5B | Alternating | 8 | E |
| MP3_2 | Serial mosquito | 3 | B | AP9_1 | Alternating | 9 | A |
| MP3_3 | Serial mosquito | 3 | C | AP9_4 | Alternating | 9 | D |
| MP3_4 | Serial mosquito | 3 | D | AP9_5 | Alternating | 9 | E |
| MP3_5 | Serial mosquito | 3 | E | AP10_1B | Alternating | 10 | A |
| MP4_1 | Serial mosquito | 4 | A | AP10_4B | Alternating | 10 | D |
| MP4_2 | Serial mosquito | 4 | B | AP10_5B | Alternating | 10 | E |
| MP4_3 | Serial mosquito | 4 | C | AP10_A | Alternating | CC | A |
| MP4_4 | Serial mosquito | 4 | D | AP10_D | Alternating | CC | D |
| MP4_5 | Serial mosquito | 4 | E | AP10_E | Alternating | CC | E |
| MP10_1 | Serial mosquito | 10 | A | BC2 | Inoculum | Stock | n/a |
| MP10_2 | Serial mosquito | 10 | B |  |  |  |  |
| MP10_3 | Serial mosquito | 10 | C |  |  |  |  |
| MP10_4 | Serial mosquito | 10 | D |  |  |  |  |
| MP10_5 | Serial mosquito | 10 | E |  |  |  |  |
| MP10_A | Serial mosquito | CC | A |  |  |  |  |
| MP10_B | Serial mosquito | CC | B |  |  |  |  |
| MP10_C | Serial mosquito | CC | C |  |  |  |  |
| MP10_D | Serial mosquito | CC | D |  |  |  |  |
| MP10_E | Serial mosquito | CC | E |  |  |  |  |

